# Scaling coalescent-based species tree inference to 100,000 taxa with STELAR-X

**DOI:** 10.1101/2025.11.22.689894

**Authors:** Anik Saha, Md. Shamsuzzoha Bayzid

## Abstract

Summary methods reconstruct species trees from collections of gene trees while accounting for gene tree discordance and provide a statistically consistent framework for phylogenomic inference under the multispecies coalescent model. While existing triplet- and quartet-based approaches such as ASTRAL and STELAR have provable statistical consistency, their running time and memory usage restrict their applicability to ultra-large datasets. We introduce STELAR-X, a statistically consistent and highly scalable triplet-based phylogenetic inference algorithm that achieves an asymptotically optimal memory complexity of *O*(*nk*) for *n* species and *k* gene trees, essentially matching the input size and allowing analyses to remain feasible as long as the input trees fit in memory, while also substantially reducing running time. STELAR-X achieves this through a compact integer tuple-based encoding of tree bipartitions, efficient precomputation of bipartition weights, and GPU parallelism. These innovations substantially reduce computational overhead in the underlying dynamic programming framework. Experiments demonstrate that STELAR-X achieves unprecedented scalability. On simulated datasets with 10,000 taxa and 1,000 gene trees, STELAR-X runs 3,576*×* faster than ASTRAL-MP (the most scalable variant of ASTRAL) while using 13.9*×* less CPU memory. STELAR-X analyzed a dataset of 100,000 taxa and 1,000 genes in 34.37 minutes using 58.40 GB RAM, and a 100,000-gene dataset with 1000 taxa in just 3.32 minutes using 74.10 GB RAM, scales that were previously intractable for statistically consistent summary methods. Moreover, applying STELAR-X to two large-scale avian datasets produced trees highly consistent with established bird phylogenies, demonstrating its robustness on biological data.

## Introduction

Widespread genome-wide discordance poses a major challenge for phylogenomic studies based on multilocus datasets (Jarvis et al. 2014, Wickett et al. 2014). A species tree represents the branching pattern of species through speciation events, while a gene tree describes the evolutionary history of a particular gene across those species. Various biological processes can cause different loci to follow different evolutionary paths, so estimating a species tree typically involves analyzing multiple genes and alignments across the genome. Among these processes, incomplete lineage sorting (ILS) is widely recognized as a dominant source of discordance, best explained under the multispecies coalescent model (Hudson 1983, Nei 1986, Nei 1987, Rosenberg 2002). In the presence of gene tree heterogeneity, traditional approaches such as concatenation–where gene alignments are merged into a single supermatrix to infer one tree–can be statistically inconsistent (Roch and Steel 2015, Degnan et al. 2009), and produce incorrect trees with high support (Kubatko and Degnan 2007).

To overcome these limitations, a range of methods have been developed that explicitly account for ILS, many of which are provably statistically consistent, meaning that they will converge in probability to the true species tree given sufficiently large numbers of genes and sites per gene (Heled and Drummond 2010, Drummond and Rambaut 2007, Liu 2008, Chifman and Kubatko 2014, Liu et al. 2010). The most scalable among these are *summary methods*, which first infer gene trees for individual loci and then combine them to estimate the species tree. Notable examples of statistically consistent, coalescent-based summary methods include MP-EST (Liu et al. 2010), ASTRAL (Mirarab et al. 2014b), BUCKy (Larget et al. 2010), GLASS (Mossel and Roch 2011), STEM (Kubatko et al. 2009), SVDquartet (Chifman and Kubatko 2014), STEAC (Liu et al. 2009), NJst (Liu and Yu 2011), ASTRID (Vachaspati and Warnow 2015), STELAR (Islam et al. 2020).

Among summary methods, ASTRAL has emerged as the most widely used and accurate approach for large-scale phylogenomic analyses under the multispecies coalescent model. By maximizing quartet consistency across input gene trees, ASTRAL provides statistically consistent species tree estimates. ASTRAL and its parallel implementation ASTRAL-MP (Yin et al. 2019) have been shown to scale to datasets with up to ∼10,000 species with several days of running time. Despite these advances, both ASTRAL and ASTRAL-MP face scalability bottlenecks when the number of species and genes exceeds several tens of thousands.

Improving the scalability of species tree inference has long been a key focus in phylogenomics. Divide-and-conquer strategies have proven particularly effective in extending summary methods to larger datasets. For example, the disk-covering method (Bayzid et al. 2014) demonstrated that partitioning taxa into overlapping subsets (“disks”), inferring local trees, and merging them can substantially enhance scalability of summary methods without sacrificing accuracy. A notable recent development in this direction is uDance (Balaban et al. 2024), a divide-and-conquer framework for inferring and incrementally updating large genome-wide phylogenetic trees. Unlike traditional phylogenomic methods that reconstruct trees *de novo* each time, uDance can update and expand an existing “backbone” tree as new genomes are added. The backbone-growth paradigm allows uDance to analyze much larger taxon sets than standard summary methods. However, constructing a “tree of life” may require building backbone trees consisting of hundreds of thousands of taxa–a challenge that remains unresolved. Furthermore, because uDance is not explicitly designed to ensure statistical consistency under the multispecies coalescent model, the scalability of provably consistent summary methods (e.g., ASTRAL) still falls far short of the needs of ultra-large phylogenomics involving hundreds of thousands of species.

In this work, we address this gap by introducing STELAR-X, an extended version of the statistically consistent, coalescent-based method STELAR (Species Tree Estimation by maximizing tripLet AgReement) (Islam et al. 2020). STELAR-X achieves unprecedented scalability in species tree inference from rooted single-copy gene trees through a combination of algorithmic redesign and efficient data representations. We introduce a new, compact integer tuple-based representation of tree bipartitions that avoids traditional *O*(*n*^2^*k*) bitset-based encoding used in widely used methods like ASTRAL. We also used a customized double-hashing scheme based on permutation-invariant and associative hash functions with provably negligible collision probability. In addition, we introduce optimized CPU and GPU parallelization to accelerate computation. These innovations yield an asymptotically optimal memory complexity of *O*(*nk*), ensuring that analyses remain feasible whenever the input gene trees fit in memory. Combined with additional algorithmic optimizations, STELAR-X reduces the running time of the original STELAR from *O*(*n*^4^*k*^3^) to *O*(*n*^2^*k*^2^) for balanced gene-trees and *O*(*n*^3^*k*^2^) for highly unbalanced cases. Through these advances, we aim to establish a scalable computational framework for statistically consistent phylogenomic inference on emerging ultra-large datasets.

## Results

### Experimental Setup

We evaluated STELAR-X on both simulated and empirical datasets. To assess scalability, we compared its performance with ASTRAL-MP (Yin et al. 2019), the only existing multi-parallel summary method, and ASTER (Zhang et al. 2025), which implements ASTRAL-IV, a faster variant of ASTRAL. For accuracy assessment, we further compared STELAR-X with ASTRAL-MP (Yin et al. 2019), wQFM-TREE (Rafi et al. 2025), ASTER (Zhang et al. 2025), and Triplet MaxCut (TMC) (Sevillya et al. 2016). All experiments were conducted on a Linux server equipped with an NVIDIA RTX 4090 GPU (24 GB VRAM) and an AMD EPYC 7742 processor with 32 active cores and 129 GB of RAM. Each analysis was allocated a maximum runtime of 48 hours.

#### Simulated Datasets

For the scalability analysis, we simulated large phylogenomic datasets under the Multispecies Coalescent (MSC) model using SimPhy (Mallo et al. 2015). In one set of experiments, we fixed the number of gene trees (*k*) at 1,000 and varied the number of taxa (*n*) from 1,000 to 100,000. In another set, we fixed *n* at 1,000 and varied *k* from 1,000 to 100,000. All datasets were generated under four model conditions corresponding to different levels of incomplete lineage sorting (ILS), ranging from low (ILS-L1) to medium-high (ILS-L4). The average gene tree discordance corresponding to these ILS levels is reported in Supplemental Table S2. To further assess the robustness of scalability, we used three more existing simulated datasets with 500-10,000 taxa (see paragraph “Scalability on estimated gene trees” of Section “Results on Scalability Analysis” in the Results). Additionally, to evaluate accuracy, we used smaller simulated datasets containing 37, 100, 200, and 500 taxa since existing methods do not scale to very large datasets. Simulation parameters are provided in Supplemental Methods 1.4 and Supplemental Table S3.

#### Biological Datasets

We analyzed two biological datasets: the avian dataset comprising 48 taxa and 14,446 genes (Jarvis et al. 2014), and an extended avian dataset comprising 363 taxa and 63,430 genes (Stiller et al. 2024).

### Results on Scalability Analysis

To assess the scalability of STELAR-X, we experimented on simulated datasets with large numbers of taxa and gene trees under medium ILS level (ILS-L2). We report the running time, CPU memory usage, and GPU VRAM consumption averaged over 5 replicates. In the first set of experiments, we fixed the number of gene trees at *k* = 1,000 and varied the number of taxa *n* from 1,000 to 100,000. Within our resource limits (128 GB CPU memory and a maximum runtime of 48 hours), ASTRAL-MP and ASTER could be executed only up to 10,000 and 7,500 taxa respectively, consistent with the scalability reported in the original studies. Consequently, we compared the methods on datasets up to these size limits (Figure 1A).

**Figure 1.**
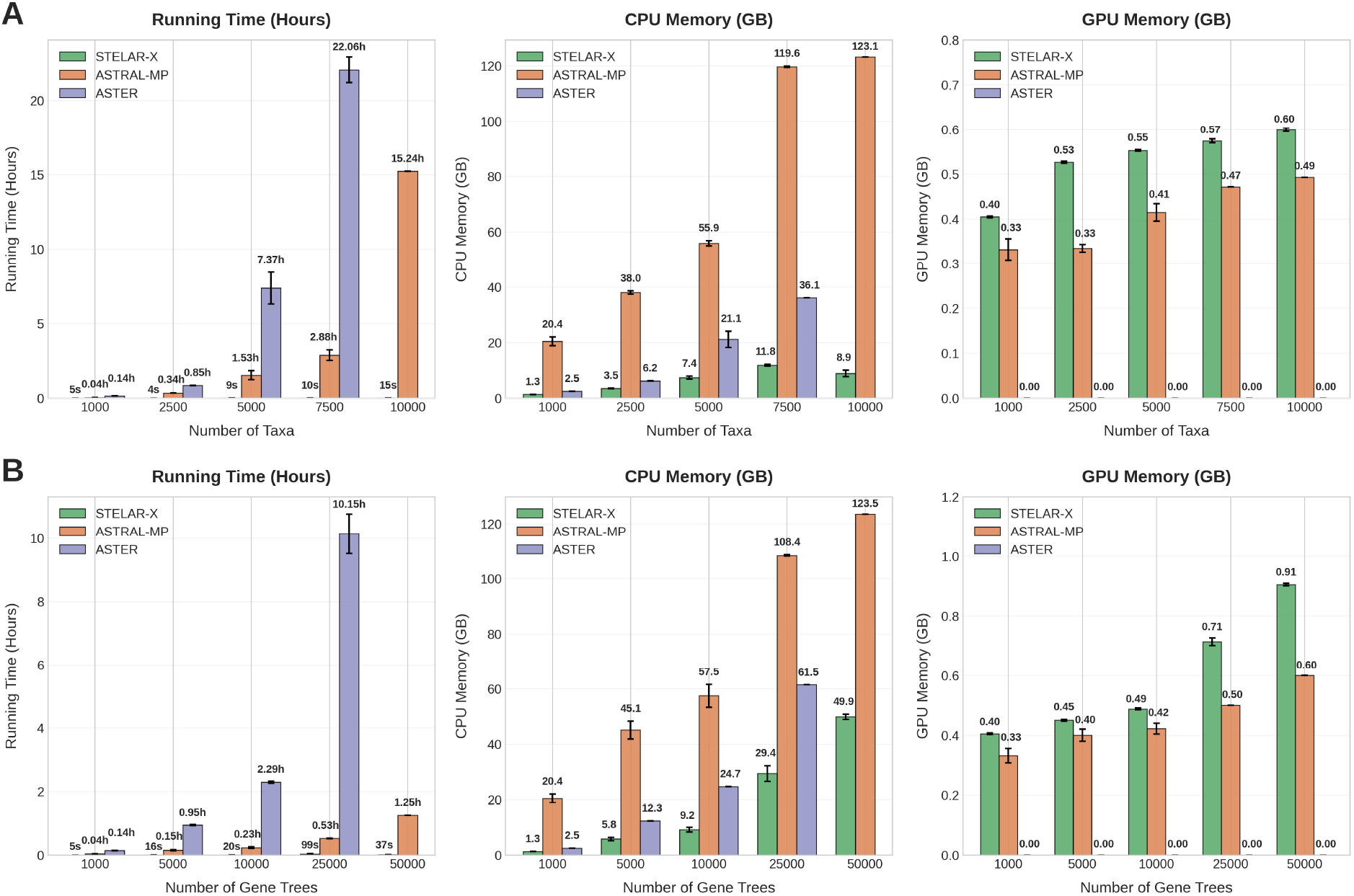
Average running time, CPU RAM, and GPU VRAM requirements of STELAR-X, ASTRALMP, and ASTER over 5 replicates (except for 10,000-taxon and 50,000-gene conditions where ASTRALMP was run on one replicate and ASTER could not be run due to their high computational demands). (*A*) 1,000 gene trees with 1,000–10,000 taxa. (*B*) 1,000 taxa and 1,000–50,000 genes.

Across all tested cases, STELAR-X consistently and substantially outperformed ASTRAL-MP and ASTER in both running time and CPU memory usage, while exhibiting a comparable trend with ASTRAL-MP in GPU VRAM utilization. In particular, ASTER cannot benefit from GPU parallelism and thus consistently took substantially longer than STELAR-X and ASTRAL-MP, yet its CPU RAM utilization was lower than that of ASTRAL-MP in all completed cases. Specifically, STELAR-X analyzed the dataset with 10,000 taxa in only 15.34 seconds, whereas ASTRAL-MP required 15.24 hours, corresponding to an approximately 3,576× speedup. STELAR-X also used only 8.87 GB of CPU RAM compared with 123.08 GB for ASTRAL-MP, an approximately 13.9× reduction in CPU memory usage. Notably, ASTRAL-MP approached the memory limit at 7,500 and 10,000 taxa, requiring 119.64 GB and 123.08 GB, respectively, whereas STELAR-X required only 11.84 GB and 8.87 GB for these two cases. As evident in Figure 1, both the running time and memory growth rates of STELAR-X are markedly lower than those of ASTRAL-MP, reflecting the asymptotically optimal *O*(*nk*) space usage achieved by our new tuple-based representation and algorithmic re-engineering. Given the observed trend, ASTRAL-MP and ASTER would require prohibitive time and memory to analyze datasets beyond 10,000 taxa, while STELAR-X remains computationally feasible well beyond this range.

Next, we evaluated scalability with respect to the number of genes while fixing the number of species at *n* = 1,000. ASTRAL-MP approached the memory limit at 50,000 genes, while ASTER exceeded the runtime limit beyond 25,000 genes. As shown in Figure 1B, STELAR-X was consistently faster and more memory-efficient across all tested dataset sizes. For example, on the 50,000-gene dataset, ASTRAL-MP required approximately 75.15 minutes and 123.54 GB of CPU RAM, whereas STELAR-X completed the analysis in only 37.2 seconds using 49.90 GB of CPU RAM. Thus, STELAR-X achieved an approximately 121× speedup while using approximately 2.48× less CPU memory. GPU VRAM usage remained modest for both methods, with STELAR-X requiring approximately 0.40–0.91 GB across these settings and showing slightly higher GPU memory requirements than ASTRAL-MP.

#### Scaling up the number of taxa and gene trees

To further evaluate our performance at a larger scale (beyond the limits of existing methods), we evaluated STELAR-X on datasets with up to 100,000 taxa and 100,000 genes. In the first experiment, we fixed *k* = 1,000 and increased *n* from 10,000 to 100,000 (Figure 2A). Consistent with our theoretical expectations, CPU memory usage exhibited an approximately linear growth, while GPU VRAM usage followed an almost perfectly linear trend. STELAR-X successfully analyzed the 100,000-taxon dataset in about 34.37 minutes using 58.40 GB of CPU RAM and 1.46 GB of GPU VRAM. For comparison, ASTRAL-MP required almost 120 GB of CPU memory to process only 7,500 taxa, and extrapolations from its growth curve suggest that analyzing 100,000 taxa would be practically infeasible for ASTRAL-MP. In contrast, given the observed linear scaling of memory and substantially faster runtime, STELAR-X is expected to handle substantially larger datasets (*>*100,000 taxa) on machines with a few hundred gigabytes of RAM (e.g., 256 GB) and modest multi-day runtimes.

**Figure 2.**
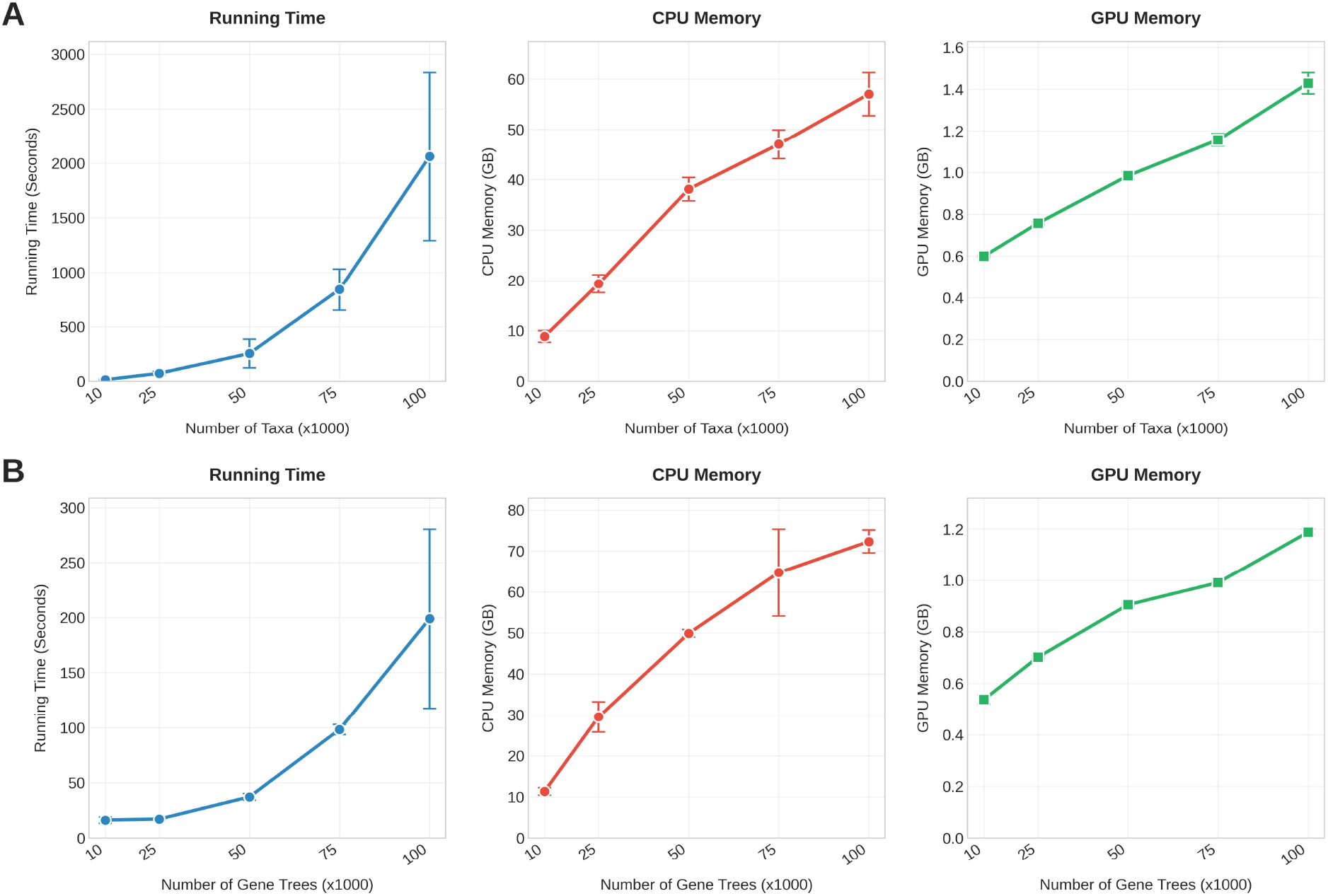
Average running time, CPU RAM, and GPU VRAM usage of STELAR-X on large datasets over 5 replicates (except for the 100,000-taxon case where we analyzed one replicate). (*A*) 10,000 to 100,000 taxa with 1,000 gene trees, (*B*) 1,000 taxa with 10,000 to 100,000 gene trees.

In the second experiment, we fixed *n* at 1,000 and varied *k* from 10,000 to 100,000 (Figure 2B). Notably, STELAR-X scaled even more efficiently with increasing numbers of genes, exhibiting nearly linear growth in runtime. It analyzed 100,000 genes in just 198.95 seconds using 74.10 GB CPU RAM– an unprecedented scale not achievable by ASTRAL-MP or other existing summary methods. CPU RAM and GPU VRAM usage remained low and continued to follow the same linear pattern.

#### Impact of varying ILS

Next, we analyzed the effect of varying levels of ILS on the running time and memory consumption of STELAR-X using simulated datasets comprising 1,000–40,000 taxa with 1,000 genes and 1,000 taxa with 1,000–50,000 genes under four model conditions, ranging from low (ILS-L1) to medium-high (ILS-L4) levels of ILS (Figure 3A-B). Consistent with our complexity analysis (see Section “Complexity Analysis” in the Methods), the running time increases with higher levels of ILS, whereas CPU RAM and GPU VRAM usage remain nearly constant, showing only minor variations due to small constant factors, thus maintaining the same linear scaling trend. For example, with 40,000 taxa, the running time increased by 3.7× from ILS-L2 to ILS-L4, while CPU RAM and GPU VRAM consumption rose only slightly, from 49 GB to 53 GB and from 0.78 GB to 0.85 GB, respectively. As the number of gene trees increased, this trend became more pronounced, with memory usage showing only negligible variation across ILS levels.

**Figure 3.**
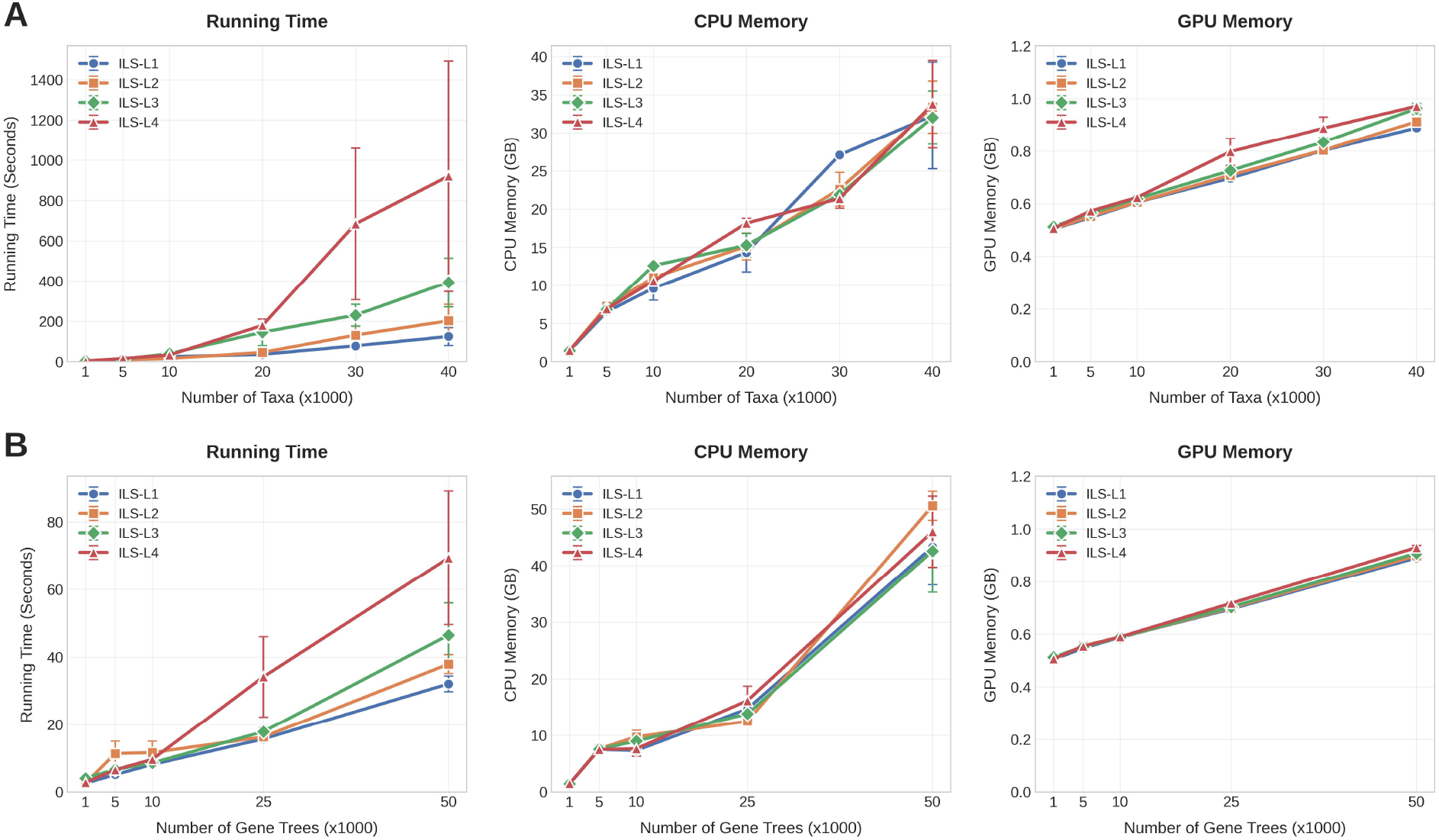
Effect of varying levels of ILS on the running time and memory usage of STELAR-X across 4 model conditions, using simulated datasets with (*A*) 1,000–40,000 taxa and 1,000 gene trees (*B*) 1,000 taxa and 1,000-50,000 gene trees.

#### Scalability on Estimated Gene Trees

To further assess the scalability of STELAR-X on estimated gene trees, we analyze three simulated datasets analyzed in prior studies. Two datasets contain 500 and 1,000 taxa, respectively, each with 1,000 genes. These datasets were previously analyzed in the ASTRAL-II (Mirarab and Warnow 2015) and later re-analyzed in the wQFM-TREE study (Rafi et al. 2025). The species trees in these datasets were generated under a Yule process with maximum tree length = 2M and speciation rate = 1 × 10^−6^. In addition, we include a larger dataset from the ASTRAL-MP study (Yin et al. 2019), comprising 10,000 taxa and 1,000 gene trees simulated with SimPhy under the multispecies coalescent model. For all three datasets, we evaluate STELAR-X and ASTRAL-MP using both the true gene trees and the corresponding estimated gene trees. For each of these datasets, the estimated gene trees were rooted using the designated outgroup taxon in that dataset. Table 1 summarizes the running time, CPU RAM usage, and GPU VRAM usage of both methods across these settings.

**Table 1.** Comparison of running time and memory usage between STELAR-X and ASTRAL-MP for true vs. estimated trees in simulated datasets.

| $n$ | $k$ | Tree type | Method | Running Time (s) | Max CPU (MB) | Max GPU (MB) |
| --- | --- | --- | --- | --- | --- | --- |
| 500 | 1000 | True | STELAR-X | 2.957 | 973.779 | 546.406 |
|  |  |  | ASTRAL-MP | 63.073 | 20434.092 | 471.501 |
|  |  | Estimated | STELAR-X | 4.173 | 1045.840 | 525.414 |
|  |  |  | ASTRAL-MP | 106.796 | 19769.439 | 448.781 |
| 1000 | 1000 | True | STELAR-X | 5.286 | 1559.008 | 536.883 |
|  |  |  | ASTRAL-MP | 232.745 | 42206.608 | 439.398 |
|  |  | Estimated | STELAR-X | 14.510 | 1885.772 | 551.219 |
|  |  |  | ASTRAL-MP | 759.608 | 42662.671 | 439.706 |
| 10000 | 1000 | True | STELAR-X | 22.430 | 10784.266 | 600.269 |
|  |  |  | ASTRAL-MP | 48950.391 | 136785.279 | 506.880 |
|  |  | Estimated | STELAR-X | 338.277 | 12275.843 | 755.200 |
|  |  |  | ASTRAL-MP | 53364.706 | 135002.436 | 860.843 |

#### Scalability under CPU-only execution

Although STELAR-X is designed as a heterogeneous CPU–GPU algorithm, we also provide a CPU-only fallback for computing environments without compatible GPU hardware. In Supplemental Results S2.1.1 and Supplemental Table S4, we compare STELAR-X with ASTRAL-MP and ASTER under CPU-only execution. STELAR-X achieves the lowest running time and CPU memory usage across all tested settings.

### Results on Accuracy of Inference

We conducted experiments on simulated datasets with 100, 200, and 500 taxa and compared the average accuracy of STELAR-X with ASTRAL-MP (Yin et al. 2019), wQFM-TREE (Rafi et al. 2025), ASTER (Zhang et al. 2025), and TMC (Sevillya et al. 2016). We could not run TMC on the 500-taxon dataset due to its high computational demand (see Supplemental Table S5). Since TMC’s accuracy was markedly lower than that of others, we report its results in the Supplemental Material (Figure S1). We also analyzed a biologically-based 37-taxon simulated mammalian dataset previously studied in (Mirarab et al. 2014a). On the 37-taxon datasets (Figures 4A,B), STELAR-X showed accuracy comparable to ASTRAL-MP, wQFM-TREE, and ASTER across varying levels of ILS and gene tree estimation error (varied by sequence length), with no statistically significant differences among methods (*p >* 0.05). STELAR-X was slightly more accurate than ASTRAL-MP on model conditions with high (0.5×) and moderate (1×) levels of ILS. wQFM-TREE achieved the highest accuracy in almost all cases. Similar patterns were observed for the 100–500-taxon datasets (Figure 4C), although STELAR-X displayed slightly higher RF rates on 100 and 200 taxa.

**Figure 4.**
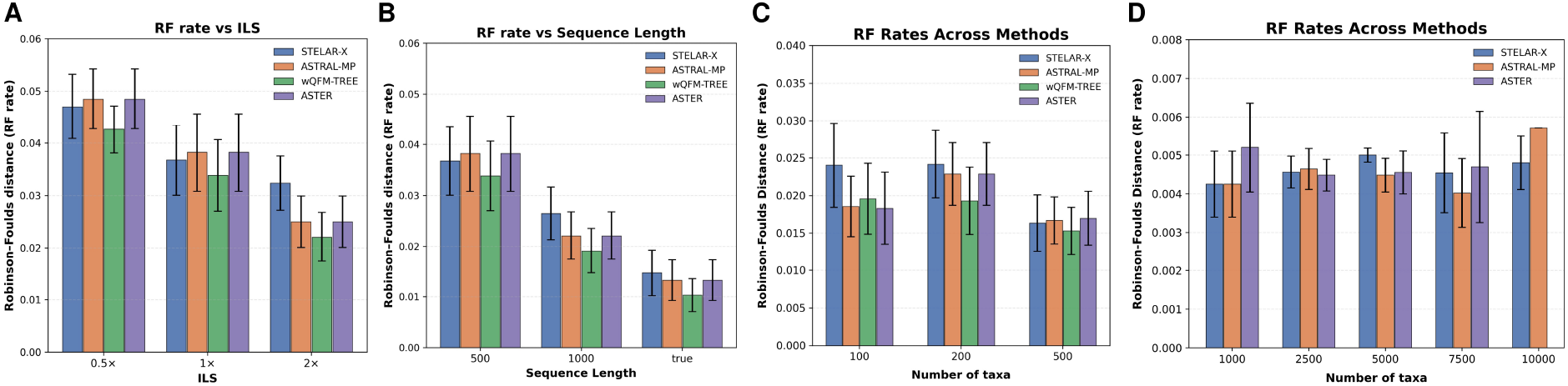
Accuracy across simulated model conditions. (*A*)-(*C*) Average RF rates of STELAR-X, ASTRAL-MP, wQFM-TREE, and ASTER over 10 replicates (*A*) varying ILS (*B*) varying sequence lengths (37 taxa) (*C*) 100–500 taxa (*D*) RF rates on 1,000–10,000 taxa over 5 replicates, except that ASTRAL-MP was run on one replicate for the 10,000-taxon condition and ASTER could not be run on the 10,000-taxon dataset.

To further assess accuracy at larger scales, we compared STELAR-X, ASTRAL-MP, ASTER on datasets spanning 1,000–10,000 taxa (excluding ASTER on 10,000 taxa due to time limit) (Figure 4D). wQFM-TREE could not be run on these datasets within our resource limits and was therefore excluded. For the 1,000-, 2,500-, and 10,000-taxon datasets, STELAR-X matched or exceeded ASTRAL-MP in accuracy, while modestly higher RF error rates were observed in the other two cases. Nonetheless, overall RF errors remained low, as the true gene trees (instead of estimated gene trees) under moderate levels of ILS were used, since sequence simulation and gene tree estimation for these large dataset sizes were computationally prohibitive.

### Results on Biological Datasets

We evaluated the avian dataset comprising 48 taxa and 14,446 genes (Jarvis et al. 2014). We rooted each gene tree in this dataset using the outgroup taxa *Struthio camelus* (ostrich) and *Tinamus guttatus* (white-throated tinamou). STELAR-X analyzed the avian dataset in only 11.36 seconds, achieving an approximately 31.5× speedup over ASTRAL-MP. We show the trees inferred by STELAR-X, ASTRAL-MP, and ASTER in Figure 5A-C. ASTER and ASTRAL-MP reconstructed an identical tree. Each of the three methods accurately reconstructed the major clades Australaves, Afroaves, Caprimulgimorphae, Core Waterbirds and Columbimorphae. However, none of the three methods could reconstruct the Cursores and Otidimorphae clades.

**Figure 5.**
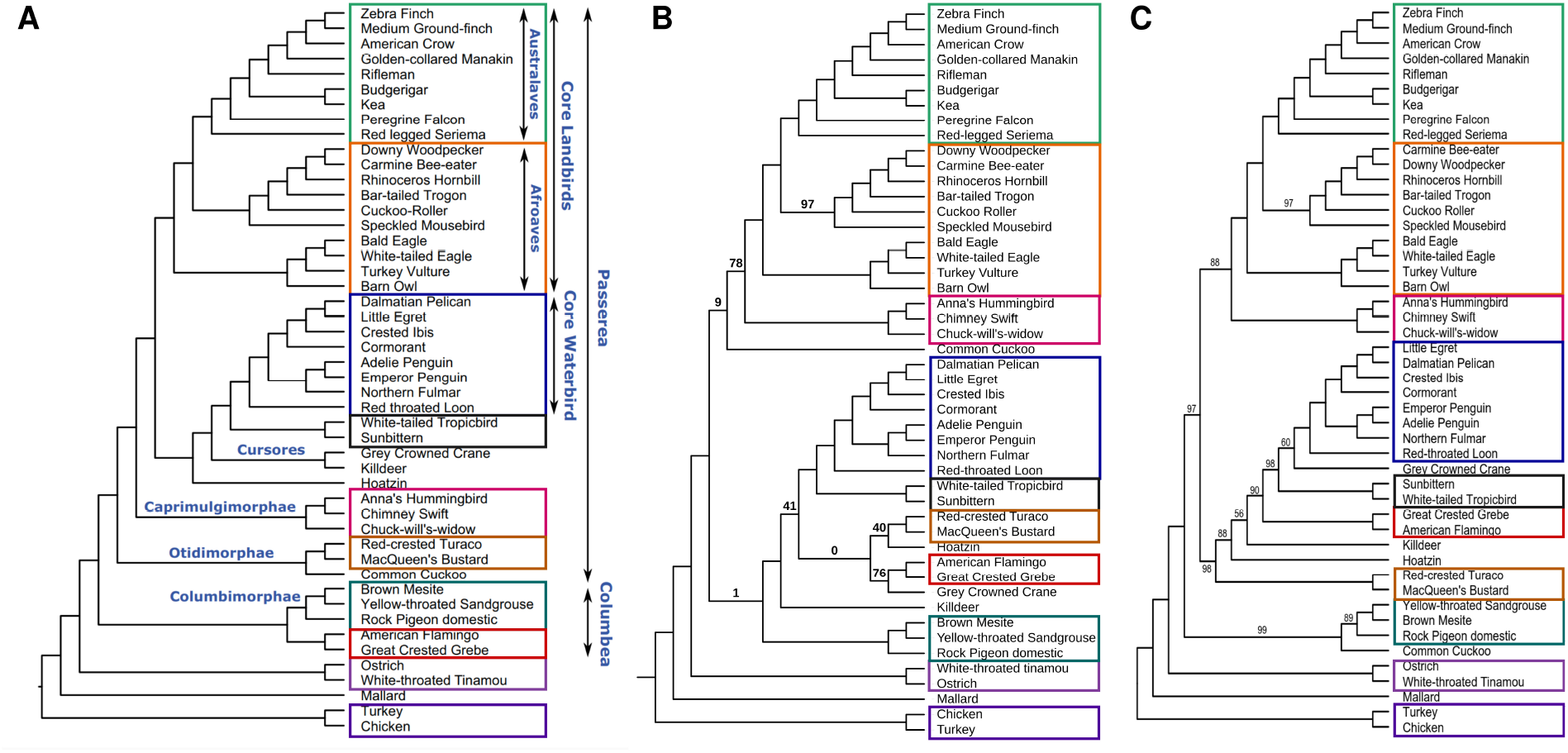
Estimated species trees for the 48-taxon avian dataset. (*A*) Species tree estimated by Binned MP-EST (published reference). (*B*) Species tree estimated by STELAR-X. (*C*) Species tree estimated by ASTRAL-MP and ASTER, which inferred topologically identical trees. The dataset comprises 48 taxa and 14,446 gene trees, with identical clades indicated. Branch support values, computed using quartet-based local posterior probabilities (Sayyari and Mirarab 2016), are shown on the branches. All branch support values are 100% except where noted.

We also analyzed an extended avian dataset comprising 363 taxa and 63,430 genes (Stiller et al. 2024). We rooted these gene trees using *Acanthisitta chloris*, which serves as an outgroup taxon (belonging to the outgroup clade Passeriformes) in that dataset. STELAR-X completed this analysis in only 5.55 minutes. It correctly reconstructed all 218 families, 37 orders, and 11 out of the 12 major clades. Figure 6 presents the species tree inferred by STELAR-X on the extended avian dataset comprising 363 taxa and 63,430 genes (intergenic loci). We also present the trees inferred by ASTRAL-MP and ASTER on this extended avian dataset in Supplemental Results 2.2 (Supplemental Figure S2). These two methods recovered identical topologies. Table 2 compares STELAR-X, ASTRAL-MP, and ASTER on the avian dataset (48 taxa, 14,446 gene trees) and the extended avian dataset (363 taxa, 63,430 gene trees) in terms of running time, CPU memory consumption, and GPU VRAM requirements.

**Table 2.** Comparison of running time and memory usage between STELAR-X, ASTRAL-MP, and ASTER on biological datasets.

| Dataset | $n$ | $k$ | Method | Running Time (s) | Max CPU (MB) | Max GPU (MB) |
| --- | --- | --- | --- | --- | --- | --- |
| avian-48 | 48 | 14446 | STELAR-X | 11.361 | 1568.988 | 547.840 |
|  |  |  | ASTRAL-MP | 357.537 | 3156.684 | 176.248 |
|  |  |  | ASTER | 143.313 | 1659.660 | 0.000 |
| avian-363 | 363 | 63430 | STELAR-X | 332.775 | 28335.602 | 825.344 |
|  |  |  | ASTRAL-MP | 42007.093 | 100356.637 | 835.584 |
|  |  |  | ASTER | 41463.458 | 53667.383 | 0.000 |

**Figure 6.**
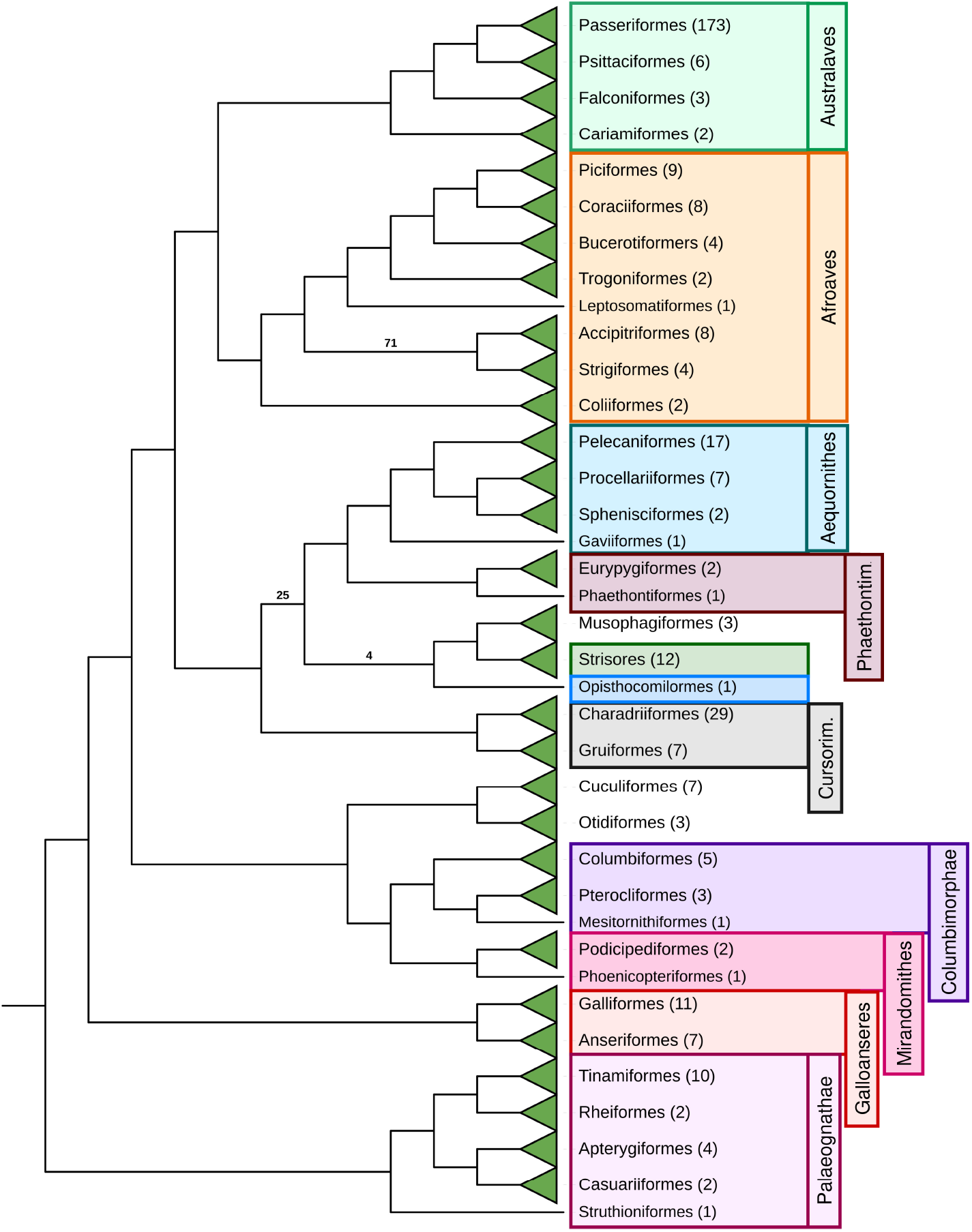
Species tree estimated by STELAR-X on the extended avian dataset with 363 taxa and 63,430 genes (intergenic loci). Although the gene trees were highly incomplete, we did not augment the set of subtree bipartitions in the gene trees by adding extra subtree-bipartitions, as our primary objective here is to demonstrate the scalability.

We now present an analysis on the reconstruction of the extended avian dataset by STELAR-X.

#### STELAR-X accurately reconstructs major phylogenetic groupings

The extended avian dataset comprises 363 taxa spanning 218 families, 37 orders, and 12 major clades (Stiller et al. 2024). STELAR-X successfully reconstructed all 218 families and all 37 orders. At the clade level, it correctly recovered 11 out of the 12 major clades, including Columbimorphae, Mirandornithes, Cursorimorphae, Aequornithes, Phaethontimorphae, Afroaves, and Australaves. The only incorrectly reconstructed clade was Otidimorphae: Musophagiformes was placed as sister to Strisores, distant from Otidiformes and Cuculiformes. Table 3 presents the reconstruction performance across the 12 clades.

**Table 3.** Recovery of major avian clades by STELAR-X on the extended avian dataset of 363 taxa.

| Clade | # Taxa | Correctly Recovered by STELAR-X |
| --- | --- | --- |
| Australaves | 184 | Yes |
| Afroaves | 38 | Yes |
| Aequornithes | 27 | Yes |
| Phaethontimorphae | 3 | Yes |
| Strisores | 12 | Yes |
| Opisthocomiformes | 1 | Yes |
| Cursorimorphae | 36 | Yes |
| Otidimorphae | 13 | No (Musophagiformes placed as sister to Strisores) |
| Columbimorphae | 9 | Yes |
| Mirandornithes | 3 | Yes |
| Galloanseres | 18 | Yes |
| Palaeognathae | 19 | Yes |

#### Mirandornithes placed as sister to Columbimorphae

While the previous study (Jarvis et al. 2014) placed Mirandornithes as sister to Columbimorphae within Columbea, the study on the 363 taxa extended avian dataset by (Stiller et al. 2024) placed Mirandornithes as sister to all other Neoaves. Interestingly, STELAR-X recovers the same previous relationship placing Mirandornithes and Columbimorphae as sisters while analyzing the extended avian dataset. They together, however are sister to all other living birds.

#### Waterbirds are placed deep within a separate clade

Unlike some previous studies (Jarvis et al. 2014, Kuhl et al. 2020) that placed waterbirds (Aequornithes and Phaethontimorphae) as sister to landbirds, the published tree corresponding to the extended avian dataset (Stiller et al. 2024) placed waterbirds deep within a diverse clade. STELAR-X also reconstructed the same relationship placing waterbirds with Strisores, Cursorimorphae, and Opisthocomiformes.

#### Australaves and Afroaves are placed as sister clades to form Telluraves

STELAR-X places Australaves and Afroaves as sisters to form Telluraves supporting the original study in (Stiller et al. 2024) and previous studies (Jarvis et al. 2014, Kuhl et al. 2020) in contrast to the trees reported in (Prum et al. 2015, Wu et al. 2024).

#### Strigiformes and Accipitriformes are placed as sisters within Afroaves

While concatenated analyses in (Stiller et al. 2024) placed Accipitriformes alone as a sister to other Afroaves, the ASTRAL-MP inferred tree reported in (Stiller et al. 2024) placed them as sister clades. STELAR-X also supported this grouping of Strigiformes and Accipitriformes within Afroaves with moderate support (71%). STELAR-X, however, places Coliiformes as sister to all other Afroaves, in contrast to (Stiller et al. 2024) which places Coliiformes deeper within Afroaves.

#### Rheiformes is grouped with Tinamiformes within Palaeognathae

In the original study on the extended avian dataset (Stiller et al. 2024), ASTRAL-MP places Rheiformes as a sister to Tinamiformes, in contrast to the study in (Cloutier et al. 2019) which placed Rheiformes as a sister to (Apterygiformes+Casuariiformes). STELAR-X also recovered the same relationship placing Rheiformes as sister to Tinamiformes within the clade Palaeognathae.

## Discussion

We presented a statistically consistent coalescent-based summary method STELAR-X, a substantially redesigned and computationally efficient extension of the triplet-based summary method STELAR. At the core of STELAR-X is a compact integer-tuple representation of subtree-bipartitions that replaces the bitset encodings used in established methods such as ASTRAL. This redesign enables extensive algorithmic optimizations and re-engineering, including efficient precomputation of bipartition weights using a customized double-hashing technique and a streamlined dynamic programming strategy with greatly reduced computational overhead. By further utilizing GPU acceleration for highly regular weight precomputations, STELAR-X delivers substantial improvements in both running time and memory efficiency.

Our experiments show that STELAR-X maintains competitive accuracy with ASTRAL-MP and wQFM-TREE across a range of simulated and biological datasets. More importantly, STELAR-X delivers unprecedented scalability, and can infer species trees from datasets comprising up to 100,000 taxa or 100,000 genes in well under an hour of computation using less than 75 GB of CPU memory, whereas current approaches encounter prohibitive memory and runtime limitations well below this scale. Notably, STELAR-X enables species tree inference with an asymptotically optimal memory complexity of *O*(*nk*), same as the input size. These results suggest that large-scale analyses can be performed without compromising statistical reliability. This capability makes it possible to apply statistically consistent inference to large-scale phylogenomic studies that were previously out of reach. In particular, STELAR-X can be used within divide-and-conquer frameworks (Bayzid et al. 2014, Balaban et al. 2024) to analyze millions of taxa. This opens the door to statistically consistent phylogenomic analyses for ambitious community initiatives to resolve the tree of life across diverse lineages, such as constructing a comprehensive tree of life for all ∼330,000 known species of flowering plants (angiosperms) (Zuntini et al. 2024, Baker et al. 2022).

Beyond scalability, the design principles introduced in STELAR-X–compact data representation, a tailored double-hashing scheme, efficient weight aggregation, and GPU-accelerated precomputation—may be applicable to other coalescent–based methods. In particular, similar ideas could be incorporated into quartet-based approaches such as ASTRAL to further improve their scalability.

This study can be extended in several important directions. First, as a triplet-based method, STELAR-X currently requires rooted gene trees. Recent work has examined how the choice of gene tree rooting strategy can affect the inference accuracy of triplet-based methods (Zaman et al. 2025). However, enabling STELAR-X to operate directly on unrooted gene trees would substantially broaden its applicability and allow a more direct comparison with widely used existing methods. Second, large biological datasets often present additional challenges, including unresolved relationships that appear as polytomies. At present, STELAR-X relies on an arbitrary resolution of such polytomies, a preprocessing step that appears to have little practical impact on species tree accuracy (Rafi et al. 2025). Nevertheless, a more principled extension would be to incorporate an explicit treatment of polytomies within the formulation. Third, it would be valuable to investigate whether augmenting the candidate set with additional bipartitions can further improve the accuracy of STELAR-X, particularly for datasets with highly incomplete gene trees. Another important direction is to extend STELAR-X to multi-copy gene family trees so that it can account for both orthology and paralogy, which are increasingly important in genome-scale datasets. Finally, applying STELAR-X to large empirical phylogenomic datasets spanning diverse lineages of the tree of life will be essential for further assessing its practical utility and robustness under realistic evolutionary conditions.

## Methods

### Problem Definition

Let *G* = {*g*_1_, *g*_2_, …, *g*_*k*_} be a collection of *k* binary rooted gene trees, each defined on the same set of *n* taxa. For any set of three taxa {*a, b, c*}, let *ab*|*c* denote the triplet tree where *a* and *b* form a cherry and *c* is the outgroup. The three possible triplet topologies for a set {*a, b, c*} of taxa are (*ab*|*c, bc*|*a, ca*|*b*). Let *τ* (*t*) represent the set of all triplets induced by a tree *t*. The problem of constructing species trees by maximizing triplet consistency with the input gene trees is defined as follows. The species tree reconstruction problem by maximizing triplet consistency seeks a species tree *T* on that maximizes the triplet score

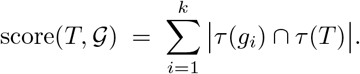

### Background: overview of STELAR

For a rooted, binary tree *t*, each internal node *u* induces a subtree bipartition (*A*|*B*) where *A* and *B* are the leaf sets of the two child subtrees of *u* (Bayzid et al. 2013). Let *SBP*(*t*) denote the set of internal bipartitions of *t*. Then the collection of subtree-bipartitions in the set *G* of gene trees is *GB* = ⋃_*Ii*_*SBP* (*g*_*i*_). We further define *UGB* as the set of unique subtree-bipartitions from *GB*. STELAR infers a species tree *ST* using a dynamic programming approach that recursively identifies the optimal (in terms of maximizing the triplet consistency) subtree-bipartitions at each level in a top-down manner (Islam et al. 2020). Note that any triplet on {*a, b, c*} is mapped to a unique internal node *u* in *T* (the least common ancestor of {*a, b, c*} in *T*), and this node in turn induces a subtree-bipartition (*A*|*B*). Consequently, the triplet consistency score of a candidate species tree *ST* can be equivalently expressed as a sum over the scores for the subtree-bipartitions in *SBP*(*ST*). Let *w*_*G*_(*x*) denote the weight of a subtree-bipartition *x* (the number of triplets in *G* that map to *x*). STELAR reconstructs the species tree by dynamically selecting subtree-bipartitions from a candidate set *CB* according to the following recursive formulation:

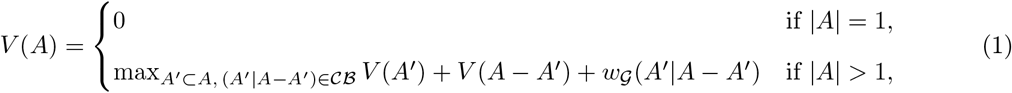

where *V* (*A*) is the optimal triplet score achievable on cluster *A* ⊆ *X*. If *CB* contains all possible bipartitions, we obtain an exact but exponential time algorithm. In the constrained version of STELAR, *CB* is restricted to only those bipartitions observed in the input gene trees, i.e., *CB* = *UGP. CB*. can also be considered as a modest extension of *UGℬ* by adding extra bipartitions for handling extensively incomplete gene trees. STELAR, including its constrained version, is a statistically consistent approach for species tree estimation (Islam et al. 2020).

### Overview of STELAR-X

To extend STELAR for the analysis of ultra-large phylogenetic datasets, we introduce algorithmic and data-structure innovations in STELAR-X: a compact tuple representation of subtree bipartitions, a permutation-invariant double-hashing scheme for frequency mapping, GPU-accelerated precomputation of bipartition weights, and an efficient dynamic-programming solver that uses the precomputed maps.

As we demonstrate theoretically (see Section “Precomputation of Bipartition Weights” in the Methods), the computation of weights constitutes the primary bottleneck in terms of running time. The structure of this computation, however, makes it particularly well-suited for Single Instruction Multiple Data (SIMD) execution (Yi 2024). We therefore isolate this step as a standalone routine and run it with GPU parallelization beforehand (see Section “Parallelization” in the Methods).

In the earlier version of STELAR, all subtree-bipartitions were represented as bitsets, the unique ones were extracted subsequently, and bitwise operations were used to calculate weights. With bitset representations, even a single taxon is represented by a bit vector of length *n*. For datasets with hundreds of thousands of taxa, this strategy leads to prohibitive RAM consumption and running time.

To address this scalability challenge, STELAR-X represents each subtree bipartition by a fixed-length integer tuple derived from the postorder leaf array of its source gene tree (see Section “Compact tuple representation of subtree bipartitions” in the Methods). It maps equivalent subtree-bipartitions (which may appear in different gene trees and in swapped order) to a unique hash signature using a customized double hashing, which makes collisions vanishingly rare, and accumulates frequencies for a subtree-bipartition in a hash table (see Section “Frequency mapping of subtree-bipartitions using permutation-invariant and associative double hashing” in the Methods).

Using this frequency map, STELAR-X precomputes the weight *w*_*G*_(*x*) for all *x* ∈ *CB* by summing contributions from all unique subtree-bipartitions. This embarrassingly parallel reduction is offloaded to the GPU (see Section “Parallelization” in the Methods). Finally, we build a cluster → subtree-bipartition map and run the DP (Equation 1) using table lookups for precomputed weights (see Section “Execution of Dynamic Programming” in the Methods). Figure 7A shows the overall algorithmic flow of STELAR-X.

**Figure 7.**
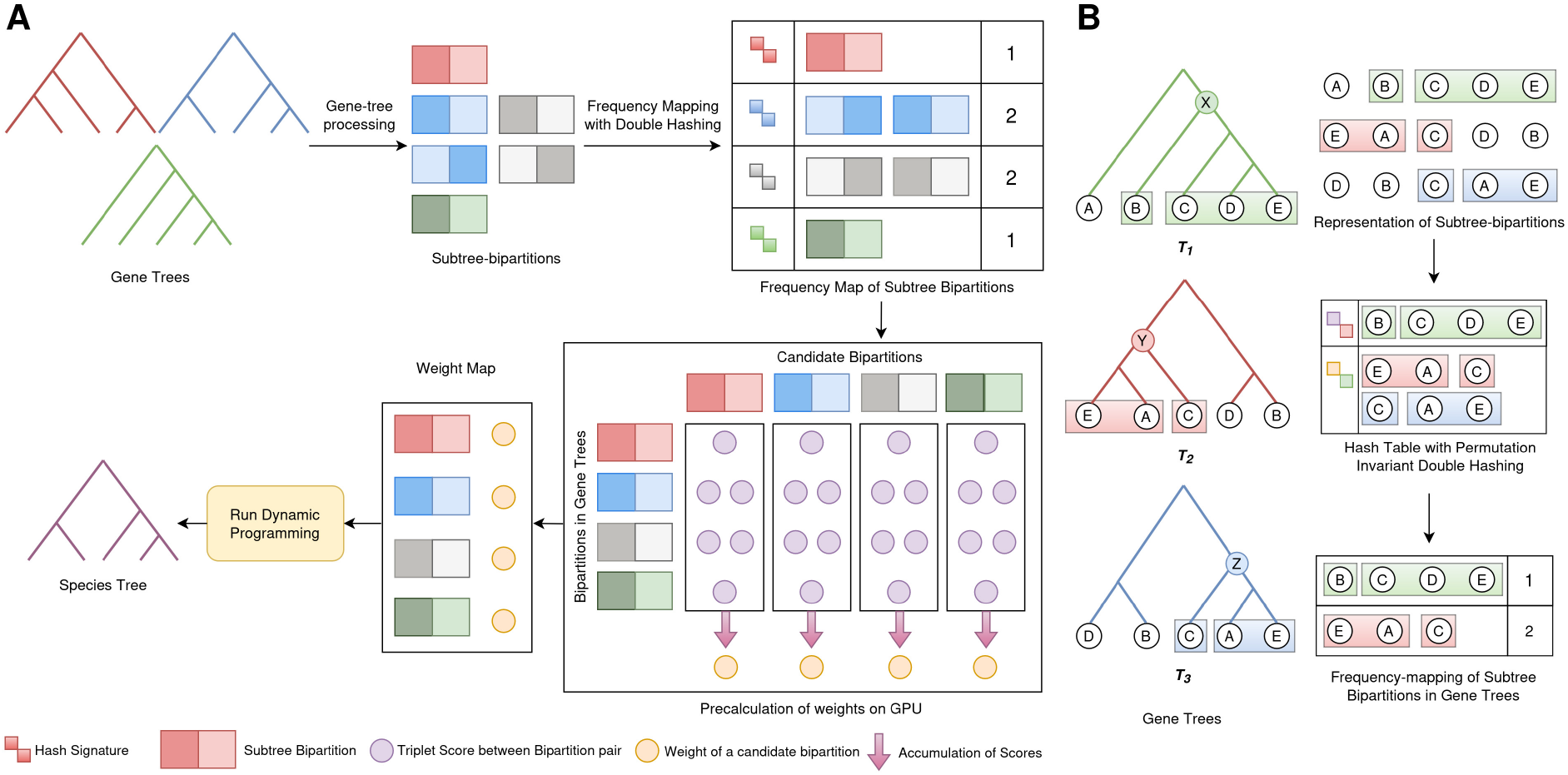
Algorithmic workflow of STELAR-X and double-hashing-based frequency mapping. (*A*) Algorithmic framework of STELAR-X integrating double-hashing-based frequency mapping, GPU-accelerated weight precomputation, and dynamic programming for species tree inference. (*B*) Illustration of frequency mapping using double hashing. Three subtree bipartitions (*B*|*CDE*), (*EA*|*C*), and (*CA*|*E*) are considered from three gene trees. Since the bipartitions (*EA*|*C*) ∈ *SBP* (*T*_2_) and (*C*|*AE*) ∈ *SBP* (*T*_3_) are equivalent, they share the same hash signature, yielding a frequency of 2.

### Compact tuple representation of subtree bipartitions

For each gene tree *g*_*i*_ we form an array *A*_*i*_ of its leaf labels in a postorder traversal. Any internal node in *g*_*i*_ induces a bipartition (*A B*) where *A* and *B*, by construction, correspond to two adjacent contiguous subarrays of *A*_*i*_. Let *A*_*i,l,r*_ denote the subarray of *A*_*i*_ from indices *l* to *r* (i.e., *A*_*i*_[*l* : *r*]). Then, if 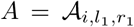 and 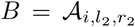, the bipartition (*A B*) can be represented as the tuple (*i, l*_1_, *r*_1_, *l*_2_, *r*_2_), where *l*_1_ ≤ *r*_1_ *< l*_2_ ≤ *r*_2_. In fact, since *l*_2_ = *r*_1_ + 1, four integers suffice to store the bipartition.

This tuple representation reduces storage from *O*(*n*) bits per bipartition to a constant number of machine words, giving an overall space of *O*(|*UGB*|) words. Because |*UGB*| = *O*(*nk*), the storage for the unique bipartitions is *O*(*nk*) words, while the usual bitset representations would take *O*(*n*^2^*k*) space. We present an example of this representation in Supplemental Methods S1.1.

### Frequency mapping of subtree-bipartitions using permutation-invariant and associative double hashing

We now describe the efficient construction of the frequency map *f*_*G*_(*x*) of a subtree-bipartition *x* ∈ *UGB* across all gene trees in G under the compact integer-tuple representation.

#### Equivalence of bipartitions

Let *P* and *Q* be sets of taxa; we write *P* ∼ *Q* if they contain the same taxa, regardless of order. Two subtree-bipartitions *x* = (*A*_1_ | *B*_1_) and *y* = (*A*_2_ | *B*_2_) are equivalent (*x* ≡ *y*) if either *A*_1_ ∼ *A*_2_ and *B*_1_ ∼ *B*_2_, or *A*_1_ ∼ *B*_2_ and *B*_1_ ∼ *A*_2_. Thus, equivalence is invariant under both taxon ordering and side swapping.

For the tuple representation of bipartitions *x* = (*i*_*x*_, *l*_1*x*_, *r*_1*x*_, *l*_2*x*_, *r*_2*x*_) and *y* = (*i*_*y*_, *l*_1*y*_, *r*_1*y*_, *l*_2*y*_, *r*_2*y*_), equivalence is possible only when *i*_*x*_ ≠ *i*_*y*_ (i.e., two bipartitions originate from different gene trees). Bipartitions within the same gene tree (*i*_*x*_ = *i*_*y*_) are either disjoint or one contains the other. A naïve cross-tree comparison to detect equivalence would require *O*(*n*) time per pair. To eliminate this overhead, we adopt a hashing-based approach that detects equivalence in *O*(1) time.

#### Permutation-invariant double hashing

Let *ϕ* be permutation-invariant and associative hash function that maps a set of taxa to a single hash value independent of order. If *x ≡ y*, by definition of equivalence, either 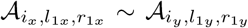 and 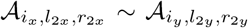, or the similar relations with the two sides swapped. Since *ϕ* is permutation-invariant, these relations imply

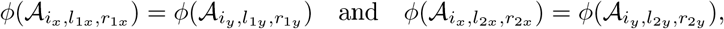

or the corresponding equalities under a side swap. Although equal hash values suggest possible equivalence, hash collisions prevent a strict guarantee. To make the probability of collisions negligible, we employ two safeguards.

First, we apply double hashing: two independent permutation-invariant and associative hash functions, *ϕ*_1_ and *ϕ*_2_, produce a composite signature (*ϕ*_1_, *ϕ*_2_) for any set of taxa. Second, we apply a per-element hash function ℋ to each taxon identifier, mapping it into 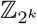 via multiplication and bitwise operations modulo 2^*k*^ so that 2^*k*^ is large. This sparse projection mitigates structural collisions common to contiguous integer labels. For any taxon *t* ∈ *X*, let id(*t*) ∈ {0, …, *n*−1} denote its identifier. For a subse *S* ⊆ *X*, we define its set-level signature as *σ*(*S*) = (*ϕ*_1_({ℋ (id(*t*)) : *t* ∈ *S*}), *ϕ*_2_({ℋ (id(*t*)) : *t* ∈ *S*})). Consequently, for a bipartition *x* = (*A* | *B*), the composite signature is (*σ*(*A*), *σ*(*B*)).

Now, since the sets *A* and *B* represent subarrays of A_*i*_, we need to efficiently compute the two values *ϕ*_1_(ℋ (id(*A*_*i,j*_)) _*l*≤*j*≤*r*_) and *ϕ*_2_(ℋ (id(*A*_*i,j*_)) _*l*≤*j*≤*r*_) for any required subarray *A*_*i,l,r*_. To this end, we precompute prefix-scan arrays for *ϕ*_1_ and *ϕ*_2_. For an associative hash function *ϕ* admitting a cancellation operator ⊖ (e.g., subtraction for addition, XOR for XOR), we define for each array A_*i*_ the prefix array *P*_*i*_ where, 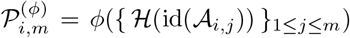 for 1 ≤ *m* ≤ *n*. Then for any subarray *A*_*i,l,r*_, the hash value of the corresponding taxon set is obtained in constant time as,

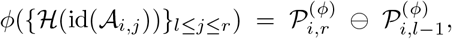

with the convention that 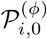 is the identity element of *ϕ*. We construct such prefix arrays independently for both *ϕ*_1_ and *ϕ*_2_, enabling *O*(1) evaluation of any required subarray hash value. Further details on the hashing scheme, along with an example hash-signature computation, are provided in Supplemental Methods S1.2 and Supplemental Table S1.

This combination of double hashing and sparse integer mapping makes collisions vanishingly rare (see the collision analysis in Theorem 1). In our implementation, *ϕ*_1_ is addition and *ϕ*_2_ is XOR over 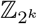, and we observed no hash collisions over extensive evaluation. All arithmetic is performed modulo 2^*k*^ so that 2^*k*^ is large (we use *k* = 64). A modulus of the form 2^*k*^ is convenient for two reasons: it preserves the associativity and group structure of bitwise operations, and it is computationally efficient because it naturally arises from the wrap-around behavior of fixed-width integer types. Finally, the frequency *f*_*G*_(*x*) of each unique bipartition *x* ∈ *UGB* is derived by inserting the computed signatures into a hash table and counting the number of bipartitions that map to the same hash value. Figure 7B schematically illustrates this process.

##### Theorem 1

(Collision Probability Bound). *Let σ*(*S*) = (*ϕ*_1_({*r*_*t*_ : *t* ∈ *S*}), *ϕ*_2_({*r*_*t*_ : *t* ∈ *S*})) *be the double-hash signature of a set S of taxa, where ϕ*_1_, *ϕ*_2_ *are independent permutation-invariant hash functions modulo M and each r*_*t*_ = ℋ (id(*t*)) *is independently and uniformly distributed in* ℤ _*M*_ . *Among B total bipartitions, the probability that at least one collision will occur is bounded by*

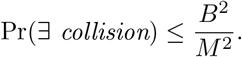

*Proof*. The key intuition is that collisions are rare because whenever two sets differ, at least one taxon contributes an independent random value, making accidental equality highly unlikely.

Let *X* be the taxon set with |*X*|= *n*. Let *B* be the number of subtree-bipartitions produced across all gene trees (in our setting, *B* = Θ(*nk*)). We choose a large modulus *M* and work in the ring ℤ_*M*_ = {0, 1, 2, … …, *M* − 1}.

#### Setup

A single-element hash

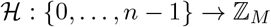

maps each taxon identifier id(*t*) to

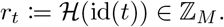

We idealize that the values {*r*_*t*_}_*t*∈*S*_ are independent and uniformly distributed in ℤ_*M*_ . We then apply two associative, permutation-invariant set-level functions *ϕ*_1_, *ϕ*_2_, each operating directly on subsets of {*r*_*t*_}. The signature of a set *S* is

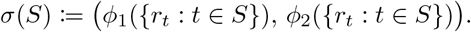

#### Single-set collision probability

Let *S* ≠ *T* and define *U* = *S \ T*, *V* = *T \ S*. Associativity ensures that *ϕ*_*a*_ admits a corresponding cancellation operator ⊖ (e.g. subtraction for addition, XOR for XOR, etc.), so that

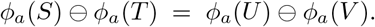

Because *U* ∪ *V* ≠ ∅, at least one *r*_*t*_ appears only in one side. Conditioning on all other *r*_*t*_, the difference reduces to a function of the uniform random variable *r*_*t*_. This equals zero with probability at most 1*/M* . Thus

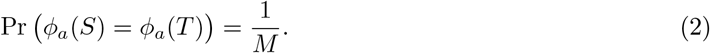

#### Pairwise signature collision

Suppose the two set-level functions *ϕ*_1_ and *ϕ*_2_ are constructed so that their collision events are statistically independent (e.g. by using independent salts or structurally distinct definitions of *ϕ*_*a*_). Then for distinct sets *S, T*,

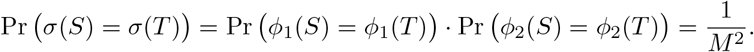

#### Collision probability for bipartitions

Let bipartitions *x* = (*A*_1_ | *B*_1_) and *y* = (*A*_2_ | *B*_2_) be distinct (non-equivalent). A collision occurs if and only if

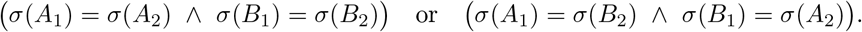

Since each matching holds with probability 1*/M* ^2^, we obtain

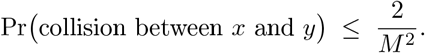

#### Global (any-pair) collision probability

Among *B* bipartitions there are at most 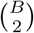 pairs. By a union bound,

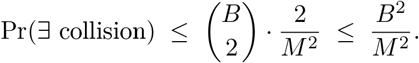

Thus, to keep this probability below a prescribed tolerance *δ*, it suffices to ensure that 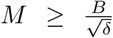. Hence, choosing *M* (*M* = 2^*k*^) sufficiently larger than *B* (*B* = Θ(*nk*)) renders global collisions negligible in practice.

### Precomputation of Bipartition Weights

We next justify and describe the precomputation of weights associated with all candidate bipartitions in *CB*. This step is essential for the dynamic programming (DP) phase of STELAR, whose state space is defined by the recursive formulation in Equation (1). The DP begins with the full taxon set *X* and recursively partitions each cluster using subtree-bipartitions. Consequently, every transition depends on the availability of subtree-bipartition weights. We now show in Theorem 2 that, in the constrained version of STELAR, this process inevitably requires the weights of all subtree-bipartitions in *CB*.

#### Theorem 2

*For the constrained version (i*.*e*., *CB* = *UGB), the DP state space of STELAR spans every bipartition x* ∈ *CB*.

*Proof*. We argue by contradiction. Suppose there exists a bipartition *x* = (*A*|*B*) ∈ *CB* that is not explored in the state space of the DP. We consider two possible cases:

#### Case 1

*A* ∪ *B* = *X* Since *CB* = *UGB*, we have *x* ∈ *GB*. As *A* ∪ *B* covers all taxa, *x* corresponds to a root-level bipartition in one of the gene trees. By construction, such bipartitions are included in the search space at the very first state transition. This contradicts our assumption that *x* is not explored.

#### Case 2

*A* ∪ *B* ≠ *X* Again, *x* ∈ *GB* and hence arises from some internal node *u* of a gene tree *g*_*i*_, where the taxa covered by *u* form the set = *A* ∪ *B*. Since ⊊ *X, u* must have a parent node *p* in *g*_*i*_. Let the bipartition induced by *p* be *y* = (*C*|*D*). By construction, either *C* = *Y* or *D* = *Y*. Thus, if *y* is explored in the DP state space, then the algorithm must also explore its child bipartition *x*, by design of the transition rules. Hence, the assumption that *x* is not explored implies that *y* is also not explored. Repeating this reasoning along the ancestor chain of *u*, we eventually reach the root of *g*_*i*_, where Case 1 applies and yields a contradiction.

Since both cases lead to contradictions, our assumption is false. Therefore, every subtree-bipartition *x* ∈ *UGB* is necessarily explored in the constrained DP state space.

Precomputing these weights therefore does not introduce redundant computation. Moreover, even when *UGB* ⊊ *CB*, precomputing weights for all *x* ∈ *CB* remains practical, provided that |*CB*| is not asymptotically larger than |*UGB*|.

#### Weight formulation

Let *x* = (*A*_1_ | *B*_1_) ∈ *CB* be a candidate bipartition and *y* = (*A*_2_|*B*_2_) ∈ *SBP* (*g*_*i*_) a bipartition from gene tree *g*_*i*_. Following (Islam et al. 2020), the number of triplets simultaneously mapped to both *x* and *y* is

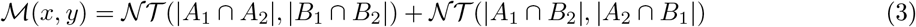

where *N T* (*n*_1_, *n*_2_) is the number of triplets mapped to a bipartition (*A* | *B*) with |*A*| = *n*_1_, |*B*| = *n*_2_:

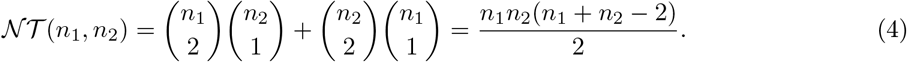

The total weight of *x* across all gene trees is then 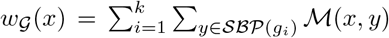. Using the precomputed frequency mapping *f*_*G*_(*y*) of unique bipartitions *y* ∈ UGB produced by the double hash table, this can be written more efficiently as,

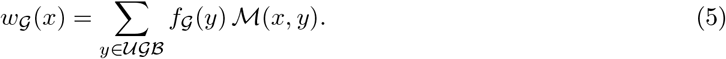

Note that each *M* (*x, y*) requires computing the four intersection cardinalities between clusters |*A*_1_ ∩ *A*_2_|, |*B*_1_ ∩ *B*_2_|, |*A*_1_ ∩ *B*_2_|, |*A*_2_ ∩ *B*_1_| (Equations (3) and (4)).

#### Efficient intersection counting using tuples

To avoid memory-intensive bitset operations, we employ our proposed integer-tuple representation of subtree-bipartitions. Let 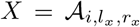 and 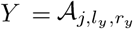 denote two clusters (*i* ≠ *j*). We have |*X*| = *r*_*x*_− *l*_*x*_ + 1 and |*Y*| = *r*_*y*_ −*l*_*y*_ + 1. The intersection |*X* ∩*Y*| is computed by iterating over each taxon in the smaller cluster and checking taxon membership in the other cluster. To optimize this membership testing, we precompute index maps *π*_*i*_(*v*), where *π*_*i*_(*v*) gives the position of taxon *v* in the post order array representation *A*_*i*_ of *g*_*i*_. Thus, for each taxon 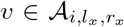, we check whether *l*_*y*_ ≤ *π*_*j*_(*v*) ≤ *r*_*y*_ (assuming |*X*| *<* |*Y* |) and accumulate the intersection count. This procedure runs in *O*(min(|*X*|, |*Y* |)) time per intersection.

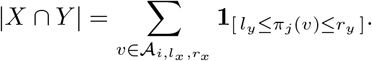

Supplemental Methods S1.1 presents an example showing the computation of *M* (*x, y*). Assuming balanced gene trees, the total time for precomputing all *w*_*G*_(*x*) over *UGB* is *O*(*n*^2^*k*^2^) (Theorem 3). Therefore, this is the dominant part of the overall running time of STELAR-X (see Section “Complexity Analysis” in the Methods). This justifies isolating the weight precomputation as a separate stage, which we further accelerate using GPU parallelization (see Section “Parallelization” in the Methods).

##### Theorem 3

*For the constrained version of STELAR (i*.*e*., *CB = UGB), if each gene tree is a perfectly balanced rooted binary tree with n* = 2^*d*^ *taxa, then the precomputation of weights requires O*(*n*^2^*k*^2^) *time*.

*Proof*. We analyze the worst-case scenario in which every bipartition across all gene trees is distinct, i.e., *UGB* = *GB*. The complexity for all other cases then follows immediately as a direct relaxation of this setting.

By Equation 5, (*w*_*G*_(*x*) = ∑_*y*∈ *UGB*_*f*_*G*_(*y*) ℳ (*x, y*)), computing the weight of each candidate bipartition requires evaluating ℳ (*x, y*) against every *y*∈ *UGB*. Thus we must evaluate ℳ (*x, y*) for every ordered pair (*x, y*) ∈ *UGB ×UGB* . We bound the total cost by first bounding the cost between two fixed balanced trees and then summing over the tree-pairs.

In a perfectly balanced rooted binary tree on *n* = 2^*d*^ leaves, an internal node at depth *ℓ* (root at depth 0) has subtree size *n/*2^*ℓ*^, and each of its two child subtrees has size *s* = *n/*2^*ℓ*+1^. Thus the set of possible child-subtree sizes is

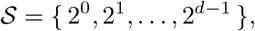

and for a given child-subtree size *s* ∈ *S* the number of internal nodes whose child-subtrees have size *s* equals the number of disjoint blocks of size 2*s* that partition the *n* leaves, i.e. 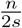. Therefore the number of subtree-bipartitions in one balanced tree whose parts (child-subtrees) have size *s* is *n/*(2*s*).

#### Per-pair cost in terms of sizes

We consider two bipartitions *x* = (*A*_1_|*B*_1_) in tree *g*_*p*_ and *y* = (*A*_2_|*B*_2_) in tree *g*_*q*_, and let the child-subtree sizes be *s* and *t*, respectively. So |*A*_1_| = |*B*_1_| = *s* and |*A*_2_| = |*B*_2_| = *t*. By Equation 3, ℳ (*x, y*) depends on four intersection counts among the parts of *x* and *y*. Each such intersection can be computed by iterating over the smaller argument using the index map *π*, hence the time to compute each intersection is *O*(min(*s, t*)). Since there are only a constant number of such intersections, the total cost to evaluate ℳ (*x, y*) is

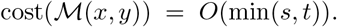

#### Cost between two balanced trees

We now group bipartitions of *g*_*p*_ and *g*_*q*_ by their child-subtree sizes *s* = 2^*i*^ and *t* = 2^*j*^ (with 0 ≤ *i, j* ≤ *d* − 1). The number of bipartitions in *g*_*p*_ with part-size *s* is *n/*(2*s*), and similarly for *g*_*q*_ with part-size *t*. Thus the number of ordered bipartition pairs with sizes (*s, t*) is 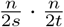, and the contribution of this size-class pair to the cost is

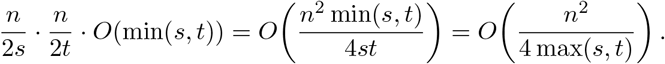

Replacing *s* = 2^*i*^ and *t* = 2^*j*^, the total cost between the two trees is

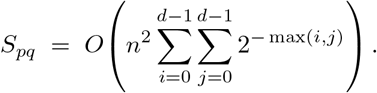

An illustration of this analysis is shown in Figure 8A for a pair of balanced binary trees of 8 taxa.

**Figure 8.**
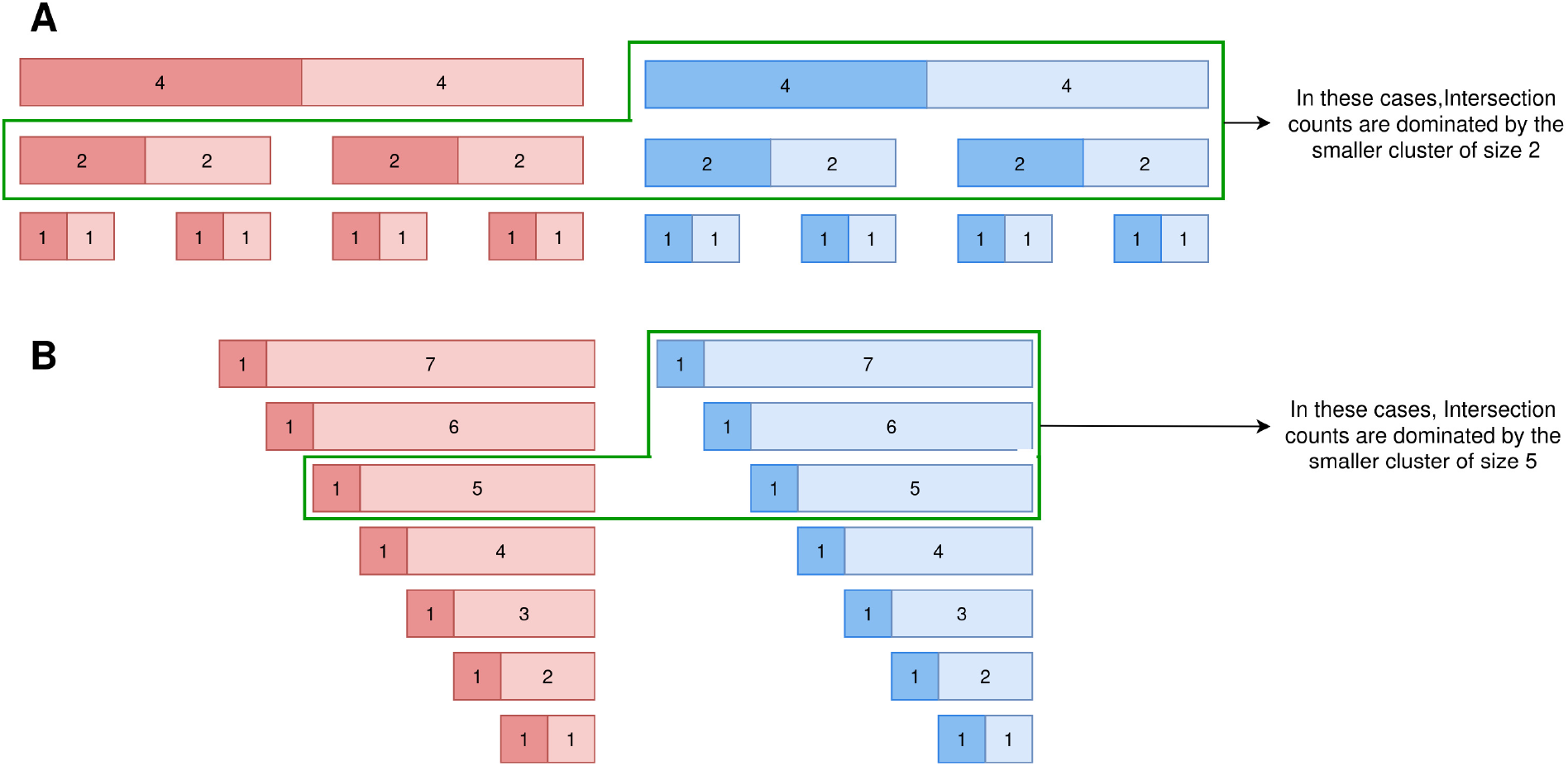
Running time analysis for computing intersection counts between two 8-taxon trees: (*A*) balanced binary trees and (*B*) caterpillar trees. Subtree bipartitions from each tree are shown, with numbers on partitions indicating the corresponding sizes.

We now evaluate the double sum by symmetry. Splitting the domain into the two triangles *i* ≤ *j* and *i > j*:

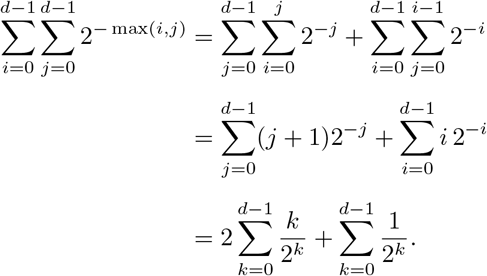

Both finite sums are uniformly bounded as *d* grows, since ∑_*k*≥0_ *k/*2^*k*^ = 2 and ∑_*k*≥0_ 1*/*2^*k*^ = 2. Hence

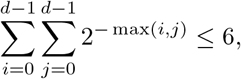

and in fact the exact value is

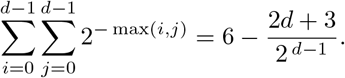

Therefore, the double sum is *O*(1). Hence *S*_*pq*_ = *O*(*n*^2^). Thus, computing all pairwise intersection costs between bipartitions of two balanced trees costs *O*(*n*^2^).

#### Extension to *k* trees

There are *k* gene trees and thus *k*^2^ ordered pairs of trees. Summing the *O*(*n*^2^) cost over all ordered pairs yields the total precomputation cost *O*(*n*^2^) *k*^2^ = *O*(*n*^2^*k*^2^), which completes the proof.

Although the analysis assumes balanced binary trees for mathematical simplicity, the practical run-time mainly depends on the smaller of the two subtrees in each bipartition pair. As a result, the observed running time typically remains close to the theoretical bound. In the extreme case of highly imbalanced fully caterpillar-like gene trees, the precomputation requires *O*(*n*^3^*k*^2^) time (Theorem 4). In practice, however, this effect is alleviated because the number of unique bipartitions |*UGB*| is generally much smaller than *nk*, making the balanced-tree assumption a good approximation of empirical behavior.

##### Theorem 4

*Assume CB = UGB. If each gene tree is a rooted caterpillar on n taxa, then the precomputation of weights (evaluating* ℳ (*x, y*) *for all ordered pairs x, y* ∈ *UGB) requires O*(*n*^3^*k*^2^) *time*.

*Proof*. The proof is provided in the Supplemental Methods S1.3, Theorem S1.1. An illustration regarding the proof is provided in Figure 8B.

### Execution of Dynamic Programming

As the final stage, we execute the dynamic programming (DP) procedure defined by Equation (1). Each DP state corresponds to a cluster of taxa, and transitions require identifying all subtree-bipartitions *x* ∈ *CB* that partition the taxa in the current cluster. To enable efficient lookup, we precompute a state-space mapping Ψ from clusters *C* ⊆ *X* to the set of candidate bipartitions.

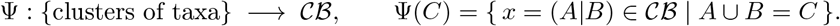

For each subtree-bipartition *x* = (*A*|*B*) ∈ *CB*, we define its associated cluster *C* = *A*∪*B* and construct a mapping. Here, again, we employ the double-hashing scheme as before and avoid the memory-intensive bitset representation. Each cluster *C* is encoded using the pair of hash signatures (*ϕ*_1_(*C*), *ϕ*_2_(*C*)). During DP (Eq. 1) we enumerate Ψ(*C*) to obtain allowed subtree-bipartitions and fetch *w*_*G*_(*A B*) in *O*(1) time per bipartition using the precomputed weight map discussed earlier (see Section “Precomputation of Bipartition Weights” in the Methods). Because both the state space (clusters) and candidate bipartitions are encoded via compact signatures, memory demand remains *O*(|*CB*| + |*UGB*|) = *O*(*nk*). The DP itself then runs in time proportional to the number of state transitions (i.e., the total number of candidate bipartitions considered), plus any bookkeeping overhead.

### Parallelization

We next identify the stages of our algorithm that are amenable to CPU and GPU parallelization and describe their implementations on multicore CPUs and GPUs. We assume *T*_*C*_ CPU threads and *T*_*G*_ GPU threads, representing the degrees of parallelism under ideal load balancing and negligible communication overhead.

#### CPU Parallelization

CPU parallelism is applied primarily to the processing of the *k* input gene trees. The collection of gene trees is divided into *T*_*C*_ approximately equal partitions, each assigned to an independent thread running in parallel. Within each thread, the parsing, subtree–bipartition extraction, and frequency mapping steps are executed sequentially on the assigned subset. When necessary, threadsafe containers and synchronization primitives are employed to ensure correct aggregation of shared data across threads.

#### GPU Parallelization

The most computationally intensive stage, computing the weight *w*_*G*_(*x*) for each candidate bipartition *x* ∈ *CB*, requires evaluating ℳ (*x, y*) for all *y* ∈ *UGB*. This corresponds to executing the same operation for *O*(|*CB*||*UGB*|) bipartition pairs, followed by accumulation along the *UGB* dimension to obtain the weights of *O*(|*CB*|) candidate bipartitions. Therefore, we offload this highly regular computation pattern to *T*_*G*_ GPU threads. We transfer the post-order leaf traversal arrays, the frequency map, and the index map to GPU. For each candidate bipartition *x* ∈ *CB*, each GPU thread computes the intersection counts for a distinct (*x, y*) pair, and results are accumulated by a reduction across *UGB* to obtain the aggregated weights. Additional optimizations, such as shared-memory tiling and coalesced access patterns, are used to reduce memory latency and improve throughput. The resulting weight map of size *O*(|*CB*|) feeds the dynamic programming algorithm.

#### Recommended Execution Mode

The weight-precomputation stage of STELAR-X exposes a very large collection of independent computations, making it particularly well suited to GPU execution, where thousands of lightweight threads can operate concurrently. Accordingly, STELAR-X is designed as a heterogeneous CPU–GPU algorithm, with the weight-precomputation stage offloaded to the GPU. A CPU-only fallback is also available for environments without compatible GPU hardware, in which case the weight-precomputation stage is parallelized across the available CPU threads. However, GPU-enabled execution is the recommended mode for fully realizing the computational benefits of STELAR-X.

### Complexity Analysis

We now provide a theoretical analysis of the running time and memory requirement of STELAR-X.

#### Theorem 5

*For n taxa, k gene trees, T*_*C*_ *CPU threads and T*_*G*_ *GPU threads, STELAR-X runs in* 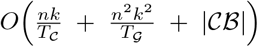 *time (under the balanced-gene-tree assumption), O*(*nk* + |*UGB*| + |*CB*|) *CPU memory, and O*(|*UGB*| + |*CB*|) *GPU memory*. *In the worst case, when all gene trees are caterpillars, the running time becomes* 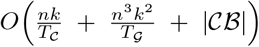.

*Proof*. We first derive bounds for the running time of STELAR-X. The parsing of the *k* input trees over *T*_C_ CPU threads requires 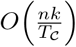 time. The subsequent computation of the frequency mapping, using the double-hashing approach, also requires 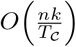 time. This efficiency follows from the use of prefix-scan arrays, which enable constant-time range hash evaluations. Next, the precomputation of weights, distributed across *T*_*G*_ GPU threads, requires 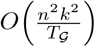 time under the balanced-gene-tree assumption as derived in Theorem 3.

Finally, we analyze the running time of the dynamic programming (DP) algorithm. By Equation 1 and Theorem 2, the size of the DP state space is at most *O*(|*CB*|). Since both the state-space map and the weight map are precomputed, the DP algorithm itself executes in *O*(|*CB*|) time.

Putting all components together, the total running time of our algorithm under the assumptions that the trees are balanced and all the bipartitions are unique is,

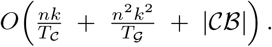

In the worst case, when all the trees are caterpillars, as derived in Theorem 4, the precomputation complexity becomes 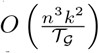, and hence the total running time becomes, 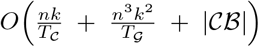.

We now analyze the memory requirements of our algorithm in both CPU RAM and GPU VRAM.

On the CPU side, storing the *k* input gene trees over *n* taxa requires *O*(*nk*) memory. The efficient representation of subtree-bipartitions, together with their subsequent processing, also requires *O*(*nk*) memory. Storing the unique subtree bipartitions in *UGB* and *CB* requires *O*(|*UGB*| + |CB|) space. The dynamic programming algorithm requires *O*(|*CB*|) additional space. Overall, the total CPU memory usage is therefore

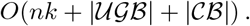

On the GPU side, memory usage is dominated by the precomputation of weights. Each bipartition is represented as a constant-length integer tuple. Consequently, the total GPU VRAM required for storing all *O*(|*UGB*| + |*CB*|) bipartitions is

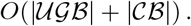

Since both |*UGB*| and |*CB*| can be bounded by *O*(*nk*) in the constrained case, the overall CPU and GPU memory consumption scales as *O*(*nk*), matching the asymptotic input-space complexity.

In practice, the precomputation complexity depends on |*UGB*| and |*CB*|. Under lower levels of ILS, |*UGB*| tends to be substantially smaller than *nk*, leading to a reduced effective running time. Consequently, the degree of ILS directly influences the practical computational cost of the algorithm.

In case of memory usage, since both |*UGB*| and |*CB*| can be bounded by *O*(*nk*) in the constrained case, the overall CPU and GPU memory usage scales as *O*(*nk*). If only the unique bipartitions were represented using traditional bitset-based representations for weight computation, the memory usage would depend on the number of distinct bipartitions, and thus on the degree of ILS. In contrast, STELAR-X employs a compact tuple-based representation throughout, achieving asymptotically optimal memory usage of *O*(*nk*) that matches the input size, regardless of the ILS level. This claim about the impact of ILS on scalability is further confirmed in our empirical results (see Section “Results on Scalability Analysis” in the Results).

### Data access

The biological datasets and their corresponding inferred species trees are available at Zenodo (https://doi.org/10.5281/zenodo.21834584).

## Code availability

The STELAR-X software, implementation, and scripts used to run the experiments are publicly available at GitHub (https://github.com/aaniksahaa/STELAR-X) and as Supplemental Code.

## Acknowledgments

This work was partially supported by the Basic Research Grant at Bangladesh University of Engineering and Technology (BUET) to MSB.

AS and MSB conceived and designed the study; AS designed and implemented STELAR-X; AS performed experiments; AS and MSB analyzed and interpreted the results; MSB supervised the study; AS wrote the first draft, and MSB and AS finalized the manuscript.

