## Supplementary Material for "Scaling coalescent-based species tree inference to 100,000 taxa with STELAR-X"

**S3 Supplementary Tables and Figures** **7**

### S1 Supplementary Methods

#### S1.1 Example: Tuple Representation of Bipartitions and Computation of $\mathcal{M}(x, y)$

We assume 1-based array indexing for this example. Consider two trees $T_1$ and $T_2$ defined over the same taxa set $\{A, B, C, D, E\}$:

$$T_1 = (A, (B, (C, (D, E)))), \qquad T_2 = (((E, A), C), (D, B)).$$

We consider the following two subtree-bipartitions ($x$ in $T_1$ and $y$ in $T_2$):

$$x = (A_1|B_1) = (B|CDE), \qquad y = (A_2|B_2) = (EA|C).$$

We first illustrate how bipartitions are represented under the post-order leaf traversal. For $T_1$, the traversal array is $\mathcal{A}_1 = \{A, B, C, D, E\}$. Let an internal node $X$ induce the subtree bipartition $x = (B|CDE)$. This bipartition $x$ is represented as the integer tuple $(1, 2, 2, 3, 5)$, where $i = 1$, $l_x = 2$, $r_x = 2$, $l_y = 3$, and $r_y = 5$. Similarly, for $T_2$, the traversal array is $\mathcal{A}_2 = \{E, A, C, D, B\}$. Let an internal node $Y$ induce the subtree bipartition $y = (EA|C)$. This bipartition is represented as the integer tuple $(2, 1, 2, 3, 3)$, where $i = 2$, $l_x = 1$, $r_x = 2$, $l_y = 3$, and $r_y = 3$.

We now compute $\mathcal{M}(x, y)$, which involves determining the sizes of intersections between the corresponding sides of these two bipartitions (Equation 3 in the main text).

$$\mathcal{M}(x, y) = \mathcal{NT}(|A_1 \cap A_2|, |B_1 \cap B_2|) + \mathcal{NT}(|A_1 \cap B_2|, |A_2 \cap B_1|)$$

To facilitate efficient comparison between bipartitions originating from different trees, we first compute the index maps $\pi_1$ and $\pi_2$. These mappings record, for each taxon, its position in the respective post-order traversal arrays $\mathcal{A}_1$ and $\mathcal{A}_2$:

$$\mathcal{A}_1 = \{A, B, C, D, E\} \;\Rightarrow\; \pi_1(A) = 1,\; \pi_1(B) = 2,\; \pi_1(C) = 3,\; \pi_1(D) = 4,\; \pi_1(E) = 5,$$
$$\mathcal{A}_2 = \{E, A, C, D, B\} \;\Rightarrow\; \pi_2(A) = 2,\; \pi_2(B) = 5,\; \pi_2(C) = 3,\; \pi_2(D) = 4,\; \pi_2(E) = 1.$$

For the intersection $A_2 \cap B_1$, we iterate over the smaller cluster $A_2 = \{E, A\}$ and test membership in $B_1 = \{C, D, E\}$ using the index maps $\pi_1$ and $\pi_2$:

$$\text{Check whether } E \in B_1: \quad \pi_1(E) = 5 \text{ and } 3 \le 5 \le 5 \;\Rightarrow\; E \in B_1 \;\Rightarrow\; \mathbf{1}_{B_1}(E) = 1,$$
$$\text{Check whether } A \in B_1: \quad \pi_1(A) = 1 \text{ and } 1 < 3 \;\Rightarrow\; A \notin B_1 \;\Rightarrow\; \mathbf{1}_{B_1}(A) = 0.$$

Summing these indicators yields

$$|A_2 \cap B_1| = \mathbf{1}_{B_1}(E) + \mathbf{1}_{B_1}(A) = 1 + 0 = 1.$$

The remaining intersections are computed similarly by iterating over the smaller of the two clusters and summing membership indicators:

$$|A_1 \cap A_2| = |\{B\} \cap \{E, A\}| = 0,$$
$$|B_1 \cap B_2| = |\{C, D, E\} \cap \{C\}| = 1,$$
$$|A_1 \cap B_2| = |\{B\} \cap \{C\}| = 0.$$

These intersection counts are then used to compute $\mathcal{M}(x, y)$.

#### S1.2 Computation of Double Hash Signatures

##### S1.2.1 Specification of Single-element Hash Function Implementation

We implement the single-element hash function $\mathcal{H}$ using a standard 64-bit multiplicative mixing scheme. For an integer taxon identifier $m$, the function applies a sequence of XOR–shift and multiplication steps:

$$\begin{aligned}
x &\leftarrow m, \\
x &\leftarrow x \oplus (x \gg 16), \\
x &\leftarrow x \times C_1, \\
x &\leftarrow x \oplus (x \gg 13), \\
x &\leftarrow x \times C_2, \\
x &\leftarrow x \oplus (x \gg 16).
\end{aligned}$$

Here $C_1 =$ `0x85ebca6b` and $C_2 =$ `0xc2b2ae35` are fixed 32-bit multiplicative constants commonly used in high-quality non-cryptographic hash functions (e.g., MurmurHash3 (Dahlgaard et al. 2017)). To avoid degenerate cases in prefix-based computations, we map the output value 0 to 1. The resulting function provides good dispersion of contiguous integer identifiers and minimizes structural collisions.

##### S1.2.2 Example: Calculation of Double Hash Signature of Bipartitions with Prefix Scan Arrays

We assume 1-based array indexing in this example. Consider the gene tree $g_i = ((D, B), (C, (A, E)))$ containing the set $\mathcal{X} = \{A, B, C, D, E\}$ of taxa. We consider a subtree-bipartition $x = (DB|CAE)$ in $g_i$, and illustrate how its double-hash signature is computed using prefix-scan arrays.

We use two hash functions $\phi_1$ and $\phi_2$, instantiated as addition and XOR, respectively. The taxa $A$–$E$ are assigned integer identifiers 0–4 sequentially. For simplicity, let the single-element hash function be defined as

$$\mathcal{H}(m) = (101m + 7) \bmod 128.$$

(see the specification of the actual single-element hash function in Section S1.2.1). We consider a small modulus $2^7 = 128$ for convenience of illustration.

The post traversal array of $g_i$ is

$$\mathcal{A}_i = \{D, B, C, A, E\}.$$

The bipartition $x = (DB|CAE)$ is therefore encoded as the integer tuple $(i, 1, 2, 3, 5)$. Prefix-sum and prefix-XOR arrays over $\mathcal{H}(\mathrm{id}(\mathcal{A}_{i,j}))$ are constructed as shown in Table S1.

We now compute the hash values for the two sides of the bipartition $x = (DB \mid CAE)$ by applying the corresponding cancellation operators— subtraction for $\phi_1$ and XOR for $\phi_2$—using the prefix-scan arrays.

**Side $\{D, B\}$ of $x$.** This corresponds to the range $\mathcal{A}_i[1:2]$. Using the prefix arrays,

$$\phi_1(\{D, B\}) = (34 - 0) \bmod 128 = 34, \qquad \phi_2(\{D, B\}) = (90 \oplus 0) \bmod 128 = 90.$$

One can verify that the sum of $\mathcal{H}(\mathrm{id}(\mathcal{A}_{i,j}))$ over the range $[1:2]$ is $(54+108) \bmod 128 = 34$. Likewise, the XOR over the same range is $(54 \oplus 108) \bmod 128 = 90$.

Thus, the signature of $\{D, B\}$ is $(34, 90)$.

**Side $\{C, A, E\}$ of $x$.** This corresponds to the range $\mathcal{A}_i[3:5]$. From the prefix arrays,

$$\phi_1(\{C, A, E\}) = (21 - 34) \bmod 128 = 115, \qquad \phi_2(\{C, A, E\}) = (23 \oplus 90) \bmod 128 = 77.$$

One can verify that the sum of $\mathcal{H}(\mathrm{id}(\mathcal{A}_{i,j}))$ over the range $[3:5]$ is $(81 + 7 + 27) \bmod 128 = 115$. Likewise, the XOR over the same range is $(81 \oplus 7 \oplus 27) \bmod 128 = 77$.

Thus, the signature of $\{C, A, E\}$ is $(115, 77)$.

**Final signature.** Combining both sides, the double-hash signature of the bipartition $x = (DB|CAE)$ is

$$((34, 90), (115, 77)).$$

#### S1.3 Theoretical analysis of the precomputation of subtree-bipartition weights

**Theorem S1.1.** *Assume $\mathcal{CB} = \mathcal{UGB}$. If each gene tree is a rooted caterpillar on $n$ taxa, then the precomputation of weights (evaluating $\mathcal{M}(x, y)$ for all ordered pairs $x, y \in \mathcal{UGB}$) requires $\mathcal{O}(n^3k^2)$ time.*

*Proof.* By Equation 5 in the main text, the precomputation requires computing $\mathcal{M}(x, y)$ for every ordered pair $(x, y) \in \mathcal{UGB} \times \mathcal{UGB}$. We first bound the cost between two fixed caterpillar trees and then sum over all ordered tree pairs.

**Bipartitions in a rooted caterpillar.** Fix a rooted caterpillar on $n$ leaves arranged along the backbone. Its internal nodes (listed from the root downwards) induce the bipartitions of sizes

$$(1 \mid n-1),\ (1 \mid n-2),\ \ldots,\ (1 \mid 1).$$

Equivalently, for each $L \in \{1, \ldots, n-1\}$ there is exactly one bipartition whose large side has size $L$ (and the small side is size 1). Thus each caterpillar has exactly $n-1$ subtree-bipartitions and the multiset of large-side sizes is $\{1, 2, \ldots, n-1\}$ (each appearing once).

**Per-pair computation of $\mathcal{M}(x, y)$.** For two bipartitions $x = (A_1 \mid B_1)$ and $y = (A_2 \mid B_2)$ the equation 3 in main text requires four intersection counts: $|A_1 \cap A_2|$, $|A_1 \cap B_2|$, $|B_1 \cap A_2|$, $|B_1 \cap B_2|$. In a rooted caterpillar at least three of these intersections involve a singleton side (size 1) and are therefore computable in $O(1)$ time. The remaining intersection is the one between the two large sides (say sizes $s$ and $t$); this intersection can be computed in $O(\min(s, t))$ time by iterating over the smaller large-side and checking membership via the index map. Thus the dominant cost for evaluating $\mathcal{M}(x, y)$ is

$$\text{cost}\left(\mathcal{M}(x, y)\right) \;=\; O\big(\min(s, t)\big),$$

where $s$ and $t$ are the sizes of the large sides of $x$ and $y$, respectively.

**Cost between two caterpillar trees.** Each of the two caterpillar trees has exactly one bipartition whose large-side size equals $L$ for each $L \in \{1, \ldots, n-1\}$. Hence the total cost between two trees equals

$$S_{pq} \;=\; \sum_{s=1}^{n-1}\sum_{t=1}^{n-1} O\big(\min(s, t)\big) \;=\; O\left(\sum_{s=1}^{n-1}\sum_{t=1}^{n-1} \min(s, t)\right).$$

An illustration of this analysis is shown in Figure 8B of main text for a pair of caterpillar trees of 8 taxa.

Now, the inner double sum is a standard combinatorial sum; counting by the minimum value $m = \min(s, t)$ gives

$$\sum_{s=1}^{n-1}\sum_{t=1}^{n-1} \min(s, t) = \sum_{m=1}^{n-1} m \cdot \#\{(s, t) : \min(s, t) = m\}.$$

For a fixed $m$, the number of ordered pairs $(s, t)$ with $\min(s, t) = m$ equals $2(n-1-m)+1 = 2n-2m-1$. Therefore

$$\begin{aligned}\sum_{s=1}^{n-1}\sum_{t=1}^{n-1} \min(s, t) &= \sum_{m=1}^{n-1} m(2n-2m-1) = 2n\sum_{m=1}^{n-1} m - 2\sum_{m=1}^{n-1} m^2 - \sum_{m=1}^{n-1} m \\ &= \Theta(n^3).\end{aligned}$$

(Using the closed forms $\sum_{m=1}^{n-1} m = \Theta(n^2)$ and $\sum_{m=1}^{n-1} m^2 = \Theta(n^3)$ one obtains the cubic bound.) Hence $S_{pq} = O(n^3)$.

**Extension to $k$ trees.** There are $k$ gene trees and thus $k^2$ ordered tree-pairs. Summing the per-pair bound $O(n^3)$ over all ordered pairs yields the total precomputation cost

$$O(n^3) \cdot k^2 = O(n^3k^2),$$

which completes the proof. □

#### S1.4 Dataset Simulation Settings

The average gene tree-gene tree (GT-GT) discordance and gene tree-species tree (GT-ST) discordance (in terms of RF rates) corresponding to four different levels of ILS are shown in Table S2. Our simulation parameters corresponding to ILS-L2 are presented in Table S3. For generating these four different ILS levels, we vary the maximum value of the uniform distribution of Effective Population Size from 150000 to 300000.

### S2 Supplementary Results

#### S2.1 Additional Results on Simulated Datasets

##### S2.1.1 Additional Results on Scalability under CPU-only execution

In Table S4, we compare the running time and CPU memory usage of STELAR-X, ASTRAL-MP, and ASTER on simulated datasets while varying the number of taxa and gene trees under CPU-only execution. STELAR-X remains the fastest and most memory-efficient method across all tested settings. In particular, ASTRAL-MP requires substantially more CPU memory than the other methods and could be run only up to 5000 taxa within our available memory, whereas ASTER exhibits substantially higher running times. On the 5000-taxon dataset with 1000 gene trees, STELAR-X was approximately 3.2× faster than ASTRAL-MP and 49.7× faster than ASTER, while using 21.3× and 3.5× less CPU memory, respectively. At 7500 taxa, where ASTRAL-MP exceeded the available memory limit, STELAR-X was 84.1× faster than ASTER while using 5.2× less CPU memory. A similar trend was observed as the number of gene trees increased. On the dataset with 25000 gene trees and 1000 taxa, STELAR-X remained slightly faster than ASTRAL-MP and 34.2× faster than ASTER, while requiring approximately 6.1× and 3.7× less CPU memory, respectively.

##### S2.1.2 Additional Results on Scalability on Simulated Datasets

We report the comparison of running time and memory usage of STELAR-X, ASTRAL-MP, ASTER, wQFM-TREE, and TMC for the simulated datasets of 37, 100, 200, and 500 taxa in Table S5. The results are averaged over 10 replicates for each setting except that we could not run TMC on the 500-taxon dataset due to its high computational demands.

##### S2.1.3 Additional Results on Accuracy on Simulated Datasets

Figure S1 presents the comparison of the accuracy of STELAR-X, ASTRAL-MP, wQFM-TREE, ASTER, and TMC averaged over 10 replicates on simulated datasets of 37, 100, 200, and 500 taxa. We could not run TMC on 500-taxon dataset due to its high computational demands.

#### S2.2 Additional Results on Biological Datasets

##### S2.2.1 Additional Results on the extended avian dataset

The species trees inferred by ASTRAL-MP and ASTER on the extended avian dataset comprising 363 taxa and 63,430 genes (intergenic loci) attained identical topologies. We report the tree in Figure S2.

#### S3 Supplementary Tables and Figures

**Table S1:** Example calculation of prefix scan arrays for two hash functions: addition and XOR on the traversal array $\mathcal{A}_i = \{D, B, C, A, E\}$. Integer identifiers 0–4 are sequentially assigned to the taxa $A$–$E$. The single element hash function $\mathcal{H}$ is assumed to be $\mathcal{H}(m) = (101m + 7) \bmod 128$. All calculations are shown modulo $2^7 = 128$.

| **j** | $\mathcal{A}_{\mathbf{i,j}}$ | id($\mathcal{A}_{\mathbf{i,j}}$) | $\mathcal{H}$(id($\mathcal{A}_{\mathbf{i,j}}$)) | **prefix sum over** $\mathcal{H}$(id($\mathcal{A}_{i,j}$)) | **prefix XOR over** $\mathcal{H}$(id($\mathcal{A}_{i,j}$)) |
|---|---|---|---|---|---|
| 0 | | | | 0 | 0 |
| 1 | D | 3 | 54 | 54 | 54 |
| 2 | B | 1 | 108 | 34 | 90 |
| 3 | C | 2 | 81 | 115 | 11 |
| 4 | A | 0 | 7 | 122 | 12 |
| 5 | E | 4 | 27 | 21 | 23 |

**Table S2:** Discordance levels under different ILS conditions analyzed in this study. We show the average gene tree-gene tree (GT-GT) and gene tree-species tree (GT-ST) discordance, measured by average RF rates, under various levels of ILS.

| **ILS Level** | **GT–GT Discordance (%)** | **GT–ST Discordance (%)** |
|---|---|---|
| ILS-L1 | 25.4 | 17.5 |
| ILS-L2 | 31.7 | 22.1 |
| ILS-L3 | 37.5 | 26.4 |
| ILS-L4 | 42.8 | 30.7 |

**Table S3:** Simulation parameters used to generate gene trees and species trees using SimPhy.

| **Parameter Name** | **Parameter Value** |
|---|---|
| Speciation rate | 0.000001 |
| Extinction rate | 0 |
| Duplication rate | 0 |
| Horizontal Gene Transfer rate | 0 |
| Ingroup divergence to the ingroup ratio | 1.0 |
| Generations | LogN(1.470055e+01,2.500000e-01) |
| Haploid effective population size | Uniform(100000,200000) |
| Global substitution rate | LogN(-1.727461e+01,6.931472e-01) |
| Species Tree Height | LogN(16.2,1) |
| Seed | 42 |

**Table S4:** Comparison of running time and CPU memory usage of STELAR-X, ASTRAL-MP, and ASTER under CPU-only execution across simulated datasets with varying numbers of taxa and gene trees. All methods were run using 32 CPU threads. Results are averaged over five replicates. ASTRAL-MP could not be run on the 7500-taxon dataset because it exceeded the available memory limit.

| $n$ | $k$ | Method | Running Time (s) | Max CPU (MB) |
|---|---|---|---|---|
| 1000 | 1000 | STELAR-X | 22.189 | 1133.322 |
| | | ASTRAL-MP | 53.199 | 22912.455 |
| | | ASTER | 511.841 | 2536.349 |
| 2500 | 1000 | STELAR-X | 215.095 | 2424.639 |
| | | ASTRAL-MP | 304.626 | 54170.608 |
| | | ASTER | 3046.142 | 6355.436 |
| 5000 | 1000 | STELAR-X | 534.318 | 6125.182 |
| | | ASTRAL-MP | 1697.536 | 130633.984 |
| | | ASTER | 26538.421 | 21639.987 |
| 7500 | 1000 | STELAR-X | 944.304 | 7065.284 |
| | | ASTRAL-MP | – | – |
| | | ASTER | 79422.350 | 36938.327 |
| 1000 | 2500 | STELAR-X | 46.271 | 2523.868 |
| | | ASTRAL-MP | 108.783 | 41536.997 |
| | | ASTER | 1932.290 | 6321.392 |
| 1000 | 5000 | STELAR-X | 98.145 | 5858.778 |
| | | ASTRAL-MP | 295.596 | 47469.815 |
| | | ASTER | 3412.275 | 12626.448 |
| 1000 | 7500 | STELAR-X | 105.701 | 7012.521 |
| | | ASTRAL-MP | 328.965 | 52675.921 |
| | | ASTER | 7316.753 | 18849.006 |
| 1000 | 10000 | STELAR-X | 125.272 | 8170.978 |
| | | ASTRAL-MP | 471.732 | 59600.042 |
| | | ASTER | 8254.236 | 25262.789 |
| 1000 | 25000 | STELAR-X | 1066.981 | 17169.490 |
| | | ASTRAL-MP | 1076.342 | 104173.445 |
| | | ASTER | 36523.173 | 62944.173 |

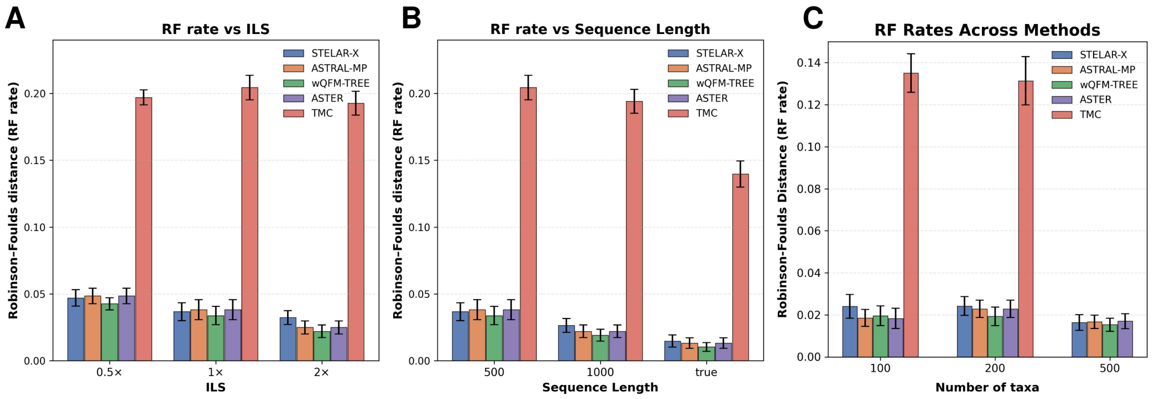

**Figure S1:** $(A)$–$(C)$ Average RF rates of STELAR-X, ASTRAL-MP, wQFM-TREE, ASTER, and TMC over 10 replicates $(A)$ varying ILS $(B)$ varying sequence lengths (37 taxa) $(C)$ 100–500 taxa (except that we could not run TMC on 500-taxon dataset due to its high computational demands.)

**Table S5:** Comparison of running time and memory usage of STELAR-X, ASTRAL-MP, ASTER, wQFM-TREE, and TMC for our simulated datasets of 37, 100, 200, and 500 taxa averaged over 10 replicates (except for the 500-taxon dataset where we could not run TMC due to its high computational demand)

| $n$ | $k$ | Method | Running Time (s) | Max CPU (MB) | Max GPU (MB) |
|---|---|---|---|---|---|
| 37 | 500 | STELAR-X | 0.692 | 224.762 | 103.465 |
| | | ASTRAL-MP | 2.36 | 403.928 | 106.196 |
| | | ASTER | 0.901 | 38.06 | 0.0 |
| | | wQFM-TREE | 1.651 | 322.547 | 0.0 |
| | | TMC | 10.689 | 146.948 | 0.0 |
| 37 | 1000 | STELAR-X | 0.683 | 219.832 | 105.677 |
| | | ASTRAL-MP | 2.485 | 422.525 | 105.472 |
| | | ASTER | 1.143 | 48.049 | 0.0 |
| | | wQFM-TREE | 2.104 | 448.374 | 0.0 |
| | | TMC | 18.71 | 190.142 | 0.0 |
| 100 | 1000 | STELAR-X | 0.884 | 444.695 | 161.587 |
| | | ASTRAL-MP | 8.411 | 1351.949 | 925.901 |
| | | ASTER | 82.461 | 279.335 | 0.0 |
| | | wQFM-TREE | 43.417 | 2170.829 | 0.0 |
| | | TMC | 927.626 | 7516.178 | 0.0 |
| 200 | 1000 | STELAR-X | 1.44 | 746.791 | 319.078 |
| | | ASTRAL-MP | 23.122 | 2590.368 | 1047.347 |
| | | ASTER | 732.518 | 505.765 | 0.0 |
| | | wQFM-TREE | 182.329 | 3729.275 | 0.0 |
| | | TMC | 15587.262 | 60137.359 | 0.0 |
| 500 | 1000 | STELAR-X | 3.697 | 1418.8 | 362.906 |
| | | ASTRAL-MP | 55.121 | 3249.421 | 789.504 |
| | | ASTER | 2979.218 | 1257.849 | 0.0 |
| | | wQFM-TREE | 1271.993 | 4788.789 | 0.0 |
| | | TMC | – | – | – |

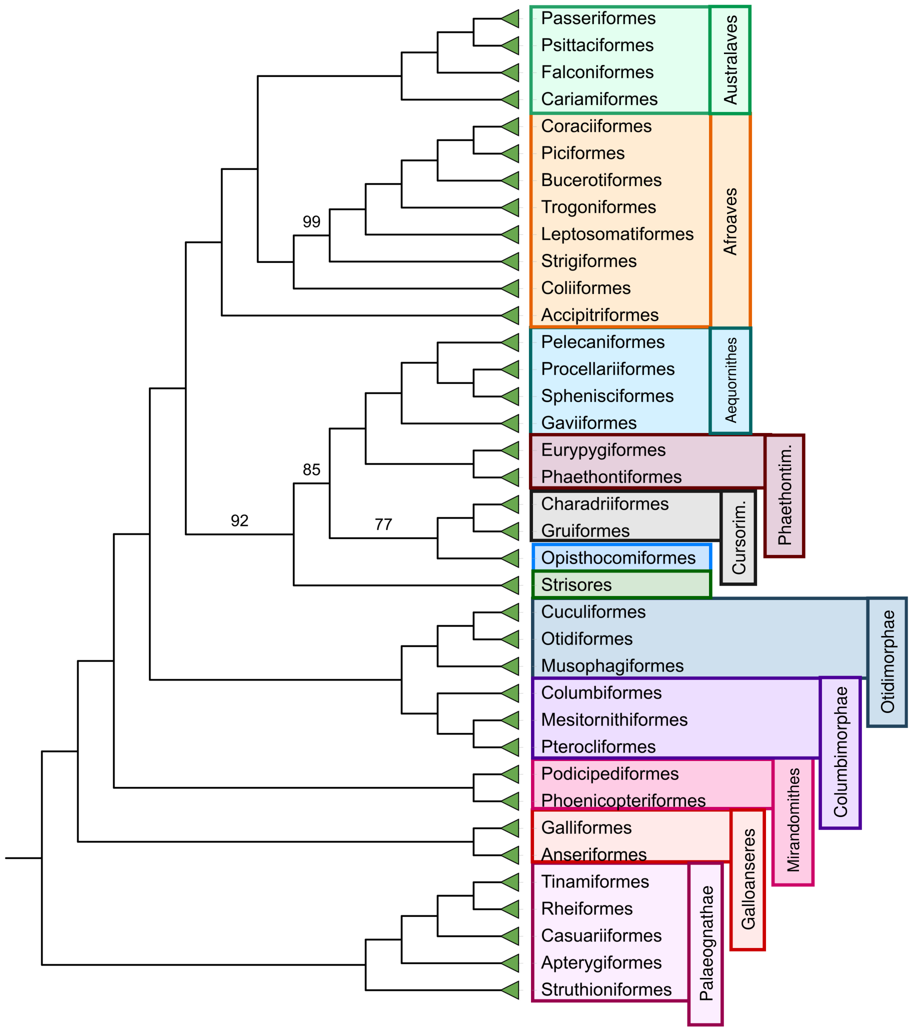

**Figure S2:** Species tree inferred independently by ASTRAL-MP and ASTER on the extended avian dataset comprising 363 taxa and 63,430 genes (intergenic loci). The two methods recovered identical tree topologies. Branch support values are 100% except where noted.
